# Association of mitochondrial fucosyltransferase TbFUT1 with the assembly of the mitochondrial F_o_F_1_-ATP synthase in bloodstream form *Trypanosoma brucei*

**DOI:** 10.1101/2023.09.01.555878

**Authors:** Samuel M. Duncan, Laura Smithson, Michele Tinti, Susan Vaughan, Michael A.J. Ferguson

## Abstract

The gene *TbFUT1* encodes an essential fucosyltransferase which, unexpectedly, localises to the mitochondrion of the protist parasite *Trypanosoma brucei*. The expression of *TbFUT1* is required for the maintenance of mitochondrial membrane potential (ΨΔm) in the bloodstream form (BSF) of the parasite, but the precise functions of TbFUT1 are unknown. Here, we demonstrate that depletion of TbFUT1 causes the accumulation of dyskinetoplastid cells; i.e., cells lacking concatenated complexes of mini- and maxicircle kinetoplast DNA (kDNA), the mitochondrial DNA of these organisms. Morphological analysis by serial face block-scanning electron microscopy showed that the dyskinetoplastid mitochondria were otherwise unperturbed with respect to structure and volume. Proteomics analyses showed that TbFUT1 depletion caused a decrease in the steady-state levels of several subunits of the F_o_-subcomplex and peripheral stalk components of the mitochondrial F_o_F_1_-ATP synthase, as well as a pronounced reduction in mitochondrial ribosomal large subunit (LSU) proteins and more minor reduction in small subunit (SSU) proteins. TbFUT1 was rendered redundant with respect to cell survival and ΨΔm generation upon F_1_-γ^WT/L262P^ mutation; a mutation that allows the generation of ΨΔm in the absence of mitochondrial translation. Additionally, depletion of TbFUT1 no longer perturbs kDNA replication in these cells, indicating that dyskinetoplasty is a downstream consequence of impaired ΨΔm. Depletion of TbFUT1 in wild type cells leads to the collapse of ΨΔm via a functional F_o_F_1_-ATP synthase complex. We therefore conclude these mutants are inhibited in the synthesis of F_o_-subcomplex components and, thus, impairing the assembly of functional F_o_F_1_-ATP synthase complexes. Curiously, mitochondrial transcript levels exhibit similar changes in abundance after FUT1 ablation in the parental and F_1_-γ^WT/L262P^ mutants. Further, the ∼5-fold overexpression of TbFUT1 in the *TbFUT1* conditional knockout mutant under permissive conditions selectively inhibits the formation of the fully RNA-edited A6 transcript by an unknown mechanism, partially suppressing F_o_F_1_-ATP synthase assembly in these mutants. Together, these data suggest that mitochondrial fucosylation is essential for the assembly of protein complexes containing kDNA encoded subunits.

## Introduction

The kinetoplastid parasite *Trypanosoma brucei* causes human African trypanosomiasis (HAT) and has a complex life cycle, with both proliferative colonising and non-proliferative transmissible lifecycle stages in the bloodstream of the mammalian host and the midgut and salivary glands of the tsetse fly vector. To establish infection and survive in these disparate environments, the parasites undergo significant changes in protein expression and metabolism (1).

The single-copy giant mitochondrion of *T. brucei* undergoes significant alterations upon parasite differentiation. The procyclic form (PCF) parasites residing in the insect midgut express a full electron transport chain able to generate ATP by oxidative phosphorylation, i.e. the concerted action of respiratory complexes I-IV, the F_o_F_1_-ATP synthase and Krebs cycle enzymes. In contrast, the mitochondrion of bloodstream form (BSF) parasites appears as a single tubule lacking the canonical respiratory chain complexes II-IV, and BSF parasites mainly generate ATP via glycolysis and maintain glycolytic NAD(H) redox by the activity of a trypanosome alternative oxidase (TAO) and glycerol-3-phosphate dehydrogenase (GPDH) (1, 2). Nevertheless, the F_o_F_1_-ATP synthase is expressed in the BSF mitochondrion where it functions in reverse mode to pump protons out of the mitochondrial matrix by hydrolysis of ATP (3, 4) to generate a mitochondrial membrane potential (ΨΔm). The latter facilitates the exchange of metabolites and import of hundreds of nuclear encoded mitochondrial proteins across the inner mitochondrial membrane. The F_o_F_1_-ATP synthase complex is composed of two subcomplexes. The matrix facing F_1_ domain functions as the catalytic unit that binds and interconverts ATP/ADP, resulting in conformational changes. The F_o_ subcomplex is composed of a c-subunit ring forming a proton pore across the inner mitochondrial membrane which is tightly associated with the highly hydrophobic mitochondrial (kDNA-encoded) A6 subunit. The F_1_ and F_o_ domains are connected by a central stalk and peripheral stator to couple conformational changes in F_1_ with rotation of the F_o_ ring.

Kinetoplastid parasites, which include the human pathogens *T. brucei, T. cruzi* and *Leishmania* spp., derive their name from the disc-shaped mitochondrial structure, the kinetoplast, that encodes their mitochondrial genes. The kinetoplast is tethered to the flagellar basal body outside of the mitochondrion. The genes encoded on the maxicircle component of kDNA include mitoribosome and respiratory chain proteins, including cytochrome reductase (CR), cytochrome oxidase (CO), NADH-dehydrogenase (ND), F_O_F_1_-ATP synthase subunits. Some kDNA-encoded transcripts must undergo RNA editing by the insertion and deletion of uridylate residues. This process is catalysed by an editosome and directed by guide RNAs (gRNAs) encoded on minicircles, the second component of kDNA (5). Wild type BSF parasites express fewer kDNA encoded genes than PCF parasites as they are inactive in oxidative phosphorylation, but the translation of mitochondrial proteins remains essential (6) due to the role of the F_O_F_1_-ATPase in maintaining ΔΨm. Thus, of kDNA-encoded genes, the A6 subunit, the small ribosomal subunit (SSU) proteins MURF5 and RPS12 and the 9S and 12S ribosomal RNAs (rRNAs) are all necessary for the translation and assembly of an active F_o_F_1_-ATP synthase in BSF parasites. The loss of kDNA (dyskinetoplasty) can be tolerated in BSF parasites by mutation of the F_1_-γ subunit, which forms part of the F_1_ subcomplex (7). These mutations have been observed in naturally akinetoplastic *T. brucei* and can be reproduced in the laboratory by reverse genetics (7). Mutation of a single leucine to a proline amino acid (L262P) presumably enables the catalytic F_1_ subcomplex to efficiently hydrolyse ATP and generate ΨΔm via ADP^3-^/ATP^4-^ exchange across the ATP/ADP inner membrane carrier (AAC) protein. By this mechanism, ΨΔm is generated in the absence of the membrane bound F_o_ pore and this confers redundancy of both kDNA replication (8, 9) and the assembly of the F_o_ subcomplex in BSF parasites. This mutation provides a useful tool to dissect the roles of enzymes involved in mitochondrial gene expression.

Essential fucosyltransferases localised to the mitochondria in *T. brucei* (TbFUT1) (10) and *Leishmania major* (LmjFUT1) (11) were recently described. Further, the ability of TbFUT1 to complement an LmFUT1 *null* mutant by the ‘plasmid shuffle’ approach suggests they have conserved functions. Both enzymes are encoded by orthologous GT11 CAZy (12) family genes and there are syntenic orthologues in all other available kinetoplastid genomes. Curiously, the ancestral *FUT1* gene appears to have been acquired from a prokaryote through horizontal gene transfer via a nucleocytoplasmic large DNA virus, rather than being inherited from an ancestral eukaryote (13). The GT11 family fucosyltransferases in other eukaryotes are generally involved in the modification of plasma membrane, secreted or lysosomal glycoproteins or glycolipids and localise to the Golgi apparatus. Therefore, both the acquisition and mitochondrial expression of TbFUT1 family members may represent a unique and significant step in kinetoplastid parasite evolution (13). Consistent with the essentiality of TbFUT1 and LmjFUT1 themselves, the *de novo* and salvage pathways of GDP-Fuc synthesis, respectively, are also essential in *T. brucei* and *L. major* (14, 15).

Despite their essentiality, little is known about the natural substrate(s) and roles of kinetoplastid mitochondrial FUT1 enzymes. However, since loss of TbFUT1 expression in BSF *T. brucei* leads to loss of mitochondrial membrane potential (ΨΔm) (10), and given that ΨΔm in BSF *T. brucei* is driven exclusively by the activity of F_o_F_1_-ATP synthase, we sought to further investigate the effects of TbFUT1 depletion on ΨΔm, mitochondrial structure and composition and on the F_o_F_1_-ATP synthase complex.

## Results

### Loss of TbFUT1 causes collapse of mitochondrial membrane potential (ΨΔm) and accumulation of dyskinetoplastic cells

Loss of TbFUT1 expression causes growth cessation and mitochondrial membrane potential collapse in BSF cells (10). These phenomena were reproduced here using a tetracycline (Tet) inducible *TbFUT1* conditional *null* mutant, also referred to here as a TbFUT1 conditional knockout mutant (TbFUT1 cKO). Thus, non-permissive (minus Tet) conditions caused cell division to cease after 2 days (Fig. 1A), with the collapse of mitochondrial membrane potential (ΨΔm) apparent in most cells by MitoTracker staining and microscopy after 3 days (Fig. 1B).

**Figure 1:**
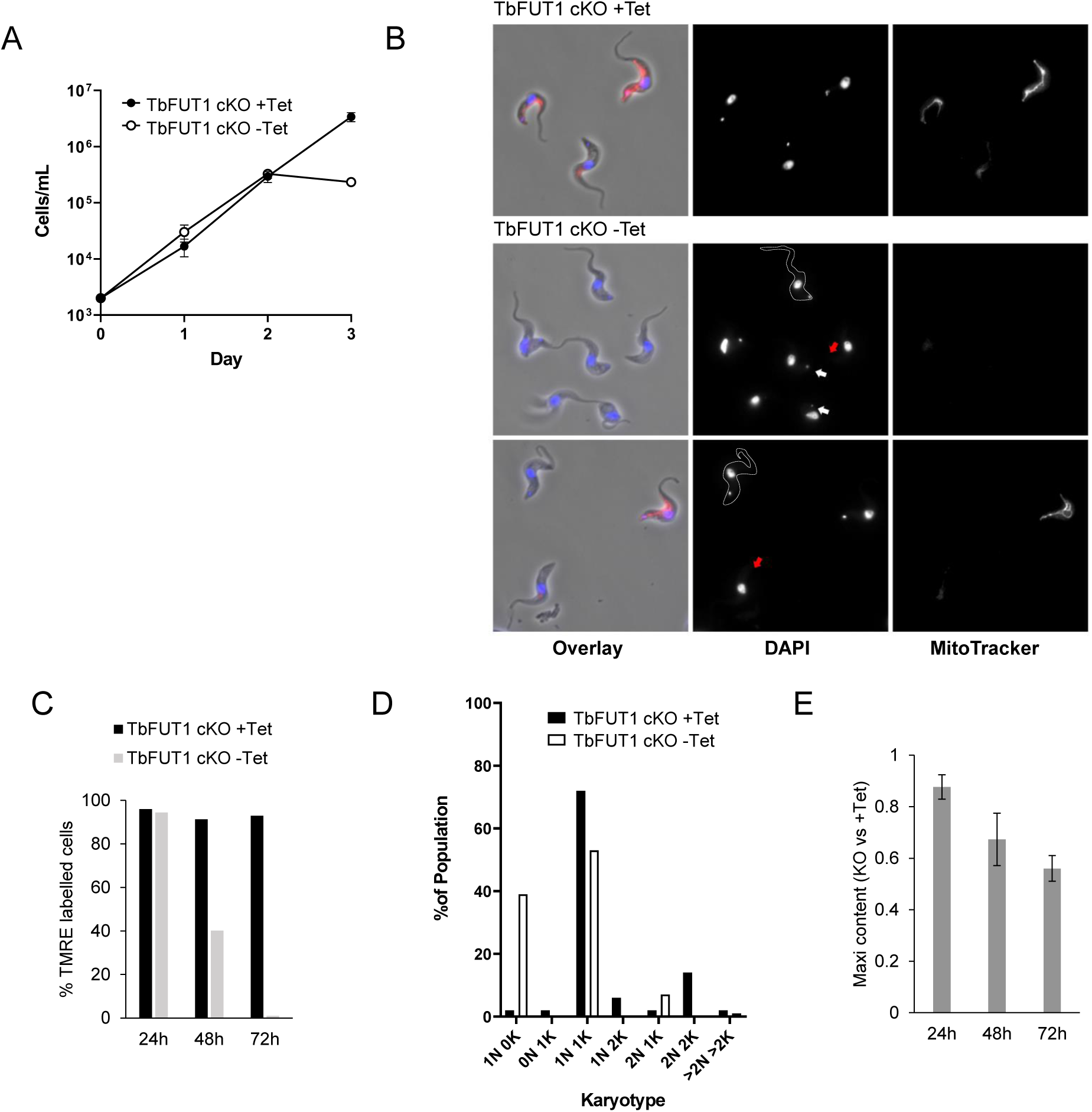
Loss of TbFUT1 causes accumulation of dyskinetoplastic cells. A. Growth of TbFUT1 cKO mutants grown in the presence (+Tet) or absence (-Tet) of Tet. BSF cells were seeded at 2×10^3^ cells/mL and counted every day for 3 days. B. Representative images showing TbFUT1 KO mutants imaged 3 days following growth in the presence (+Tet) or absence (-Tet) of tetracycline. Live cultures were treated with 10 nM MitoTracker for 30 mins prior to fixation, and the dye is shown in red merged with DNA stained by DAPI in blue in the overlay panel. Red arrows indicate cells lacking any detectable kDNA, white arrows indicate cells with kDNA of reduced size cells lacking MitoTracker uptake but retaining normal kDNA staining are outlined. C. Flow cytometry analysis of live WT and TbFUT1 cKO mutants cells treated with TMRE after growth in the presence (+Tet) or absence (-Tet) of tetracycline for 24, 48 and 72 hours. A non-stained WT control was included as a negative control and a TMRE positive gate drawn based on wild type fluorescence levels. D. Karyotype analysis of TbFUT1 cKO mutants grown in the presence or absence of tetracycline for 3 days. Counts are representative of approximately 110 cells per treatment. E. Southern blot detection of Maxicircle kDNA harvested from cells 24, 48 and 72 hours following TbFUT1 KO (-Tet). Quantification of Maxicircle DNA relative to the loading control was calculated using ImageJ gel peak analysis software. The normalised expression values were used to compare band intensity between TbFUT1 cKO cells ± Tet.

Quantitative and time-resolved analysis, using the membrane potential dependent dye tetramethyl rhodamine ethyl ester (TMRE) and live cell flow cytometry, showed that ΨΔm was unaffected 24 h after Tet removal but was significantly reduced and virtually ablated 48 h and 72 h after Tet removal, respectively (Fig. 1C, Fig. S1C).

In addition, we noted by microscopy that cells grown under non-permissive conditions for 3 days were heterogeneous with respect to kinetoplast kDNA staining with DAPI: some cells appeared normal, whereas others appeared to either lack (i.e., dyskinetoplastic cells) or contain less kDNA (Fig. 1B, see cells with red and white arrows, respectively). Effects on kDNA content were also apparent after 2 days of culture without Tet (Fig. S1A). Some cells were observed lacking MitoTracker uptake but retaining normal kDNA staining (Fig. 1B, outlined cells).

Quantitative karyotype analysis by DAPI staining and fluorescent microscopy of the TbFUT1 cKO mutant grown in the presence or absence of tetracycline for 3 days indicated that ∼40% of cells become dyskinetoplastic (1N0K) under non-permissive conditions (Fig. 1D). Southern blotting was performed using probes to maxicircle kDNA and to a nuclear hexose transporter gene loading control (Fig. 1E, Fig. S1B). The TbFUT1 cKO cells grown under -Tet conditions for 2 or 3 days had ∼35% and ∼45% less maxicircle DNA content, respectively, relative to the cells grown in +Tet conditions.

Taken together, we suggest that ΨΔm collapse under non-permissive condition in the TbFUT1 cKO cell line precedes dyskinetoplasty and growth arrest.

### Serial block face scanning electron microscopy (SBF-EM) analysis of dyskinetoplasty caused by loss of TbFUT1

To further investigate the onset of dyskinetoplasty, we performed serial block face scanning electron microscopy (SBF-EM) analysis on TbFUT1 cKO cells harvested after 77 h of growth in the presence or absence of Tet. Whereas the +Tet cells showed normal rod-like kDNA (Fig. 2A, white arrow) within typical kinetoplast bulge structures (Fig. 2B, white arrows), the -Tet cells contained more amorphous material inside the otherwise apparently unaffected kinetoplast bulges (Fig 2A and B, red arrows). The presence of residual DNA in the bulges of the -Tet TbFUT1 deficient cells was demonstrated by longer exposure of the DAPI stained mutants, revealing low levels of fluorescent signal emanating from the kDNA region (Fig. S2). These data suggested a deficiency in the replication of kDNA. The SBF-EM analysis of different cell cycle stages identified a high proportion (∼80%) of cells containing disordered kDNA after 77 h without Tet (Table S1). These data indicate that perturbations to kDNA structure occur upon TbFUT1 depletion.

**Figure 2:**
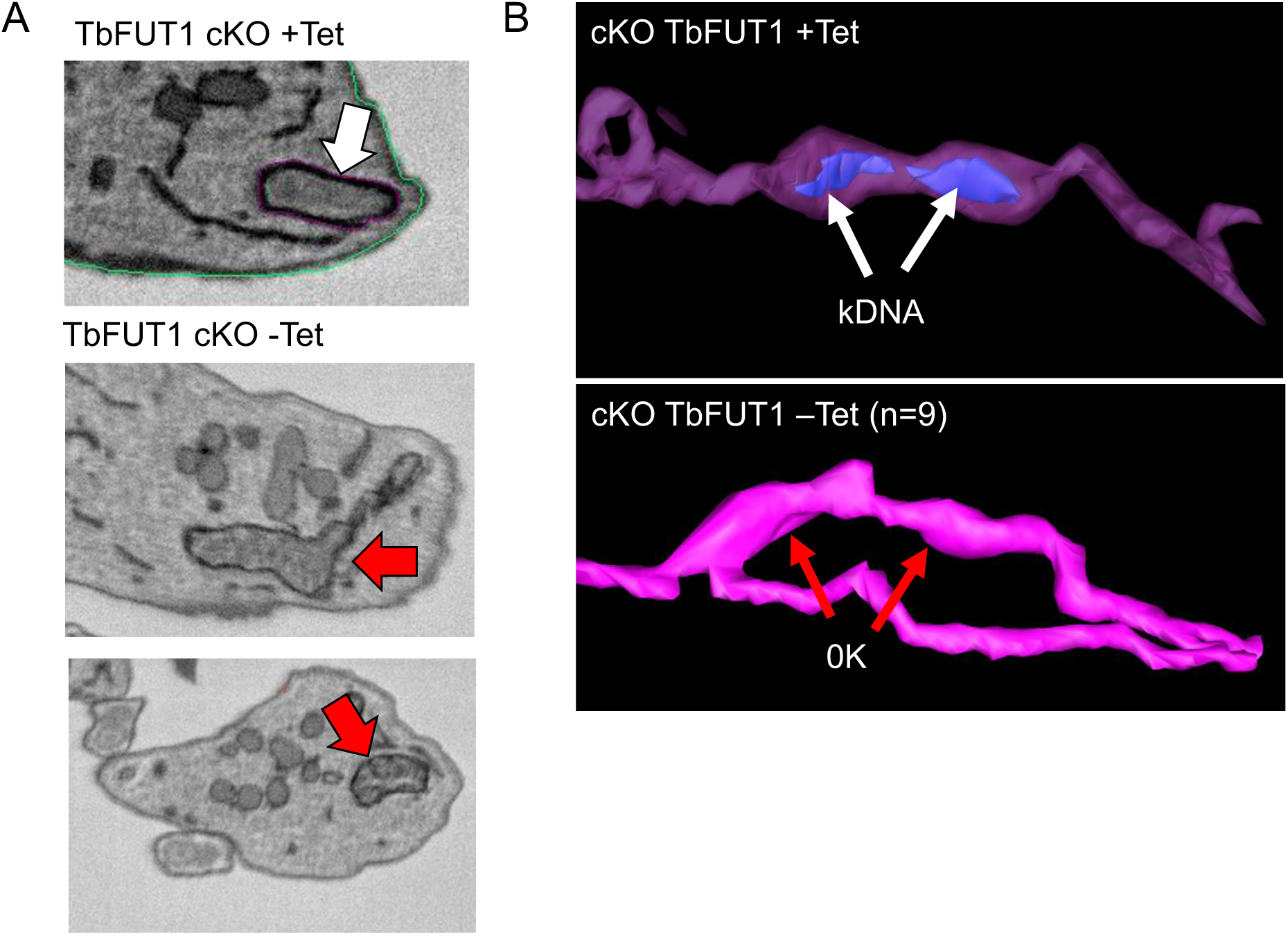
Loss of TbFUT1 has no observable effect on mitochondrial structure. WT and TbFUT1 cKO mutants cells were analysed after growth in the presence or absence of tetracycline for 3 days. A. Representative images of SBF-EM analysed cells stained to enhance membrane detection. The rod-shaped kDNA residing within the membranous ‘bulge’ is evident in TbFUT1 expressing cells (+Tet, white arrow) but the kDNA within the bulge becomes fragmented and disordered upon TbFUT1 loss (-Tet, red arrows). B. SBF-EM analysis of the mitochondria from 2K1N TbFUT1 cKO mutants was used to generate 3D models representing the mitochondrial structure. In +Tet conditions the kDNA residing within the pocket is shown in blue (white arrows). Upon loss of TbFUT1 the mitochondrial DNA is no longer detected after imaging 9 separate cells (-Tet, red arrows).

The SBF-EM analysis was used to further explore mitochondrial structure in the TbFUT1 cKO cell line grown for 3 days ±Tet. Nine individual cells having undergone kDNA division and ready to undergo nuclear division (i.e., 1N2K cells), were selected for this analysis. However, despite -Tet cells exhibiting disordered kDNA, there were no observable differences in mitochondrial morphology or ultrastructure between these cells (Fig. 2B). Finally, the mitochondrial volume was calculated by analysis of eight individual 1N2K cells from each treatment, revealing no change in mitochondrial volume following TbFUT1 depletion (Table 1).

**Table 1.**
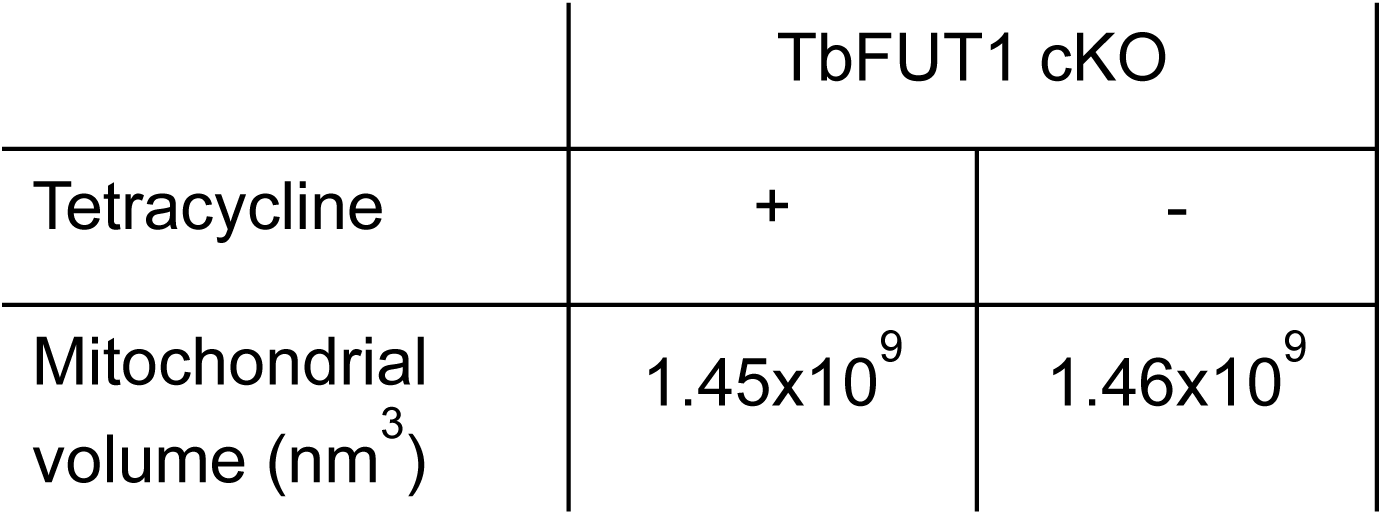
Mitochondrial volume is not affected by TbFUT1 KO. Average mitochondrial volume of 2K1N TbFUT1 cKO cells grown in the presence or absence of tetracycline for 77h (n=8).

Taken together, these data suggest that loss of TbFUT1 expression in BSF *T. brucei* leads to specific effects on mitochondrial ultrastructure relating to kDNA content.

### TbFUT1 is not essential for kDNA maintenance

From the experiments described above, TbFUT1 appears to be involved in some way in kDNA replication and/or segregation and/or stability. However, it was not clear whether TbFUT1 is directly *required* for kDNA maintenance, or if the observed phenotype was a secondary consequence. As described in the introduction, an L262P mutation of the F_1_-γ subunit of the F_o_F_1_-ATP synthase complex allows BSF *T. brucei* to survive in the absence of kDNA (7). Consequently, in an L262P mutant background the downregulation of genes necessary for kDNA synthesis and maintenance generally leads to the spontaneous loss of the mitochondrial kDNA without affecting cell growth (8, 9).

Thus, to investigate whether TbFUT1 is involved directly in kDNA maintenance, we modified our TbFUT1 cKO mutant by conferring it with single allele (TbFUT1 cKO/F1-γ^WT/L262P^) and double allele (TbFUT1 cKO/F1-γ^L262P/L262P^) point mutations in F_1_-γ (Fig. S3). The resulting mutants retained Tet inducible TbFUT1 expression (Fig. 3A) and continued to maintain their kDNA and ΔΨ_m_ dependent MitoTracker staining, despite depletion of TbFUT1 in -Tet conditions (Fig. 3B). As expected, these same cells, nevertheless, remained viable after induction of an akinetoplastid state by treatment with ethidium bromide (Fig. 3C).

**Figure 3:**
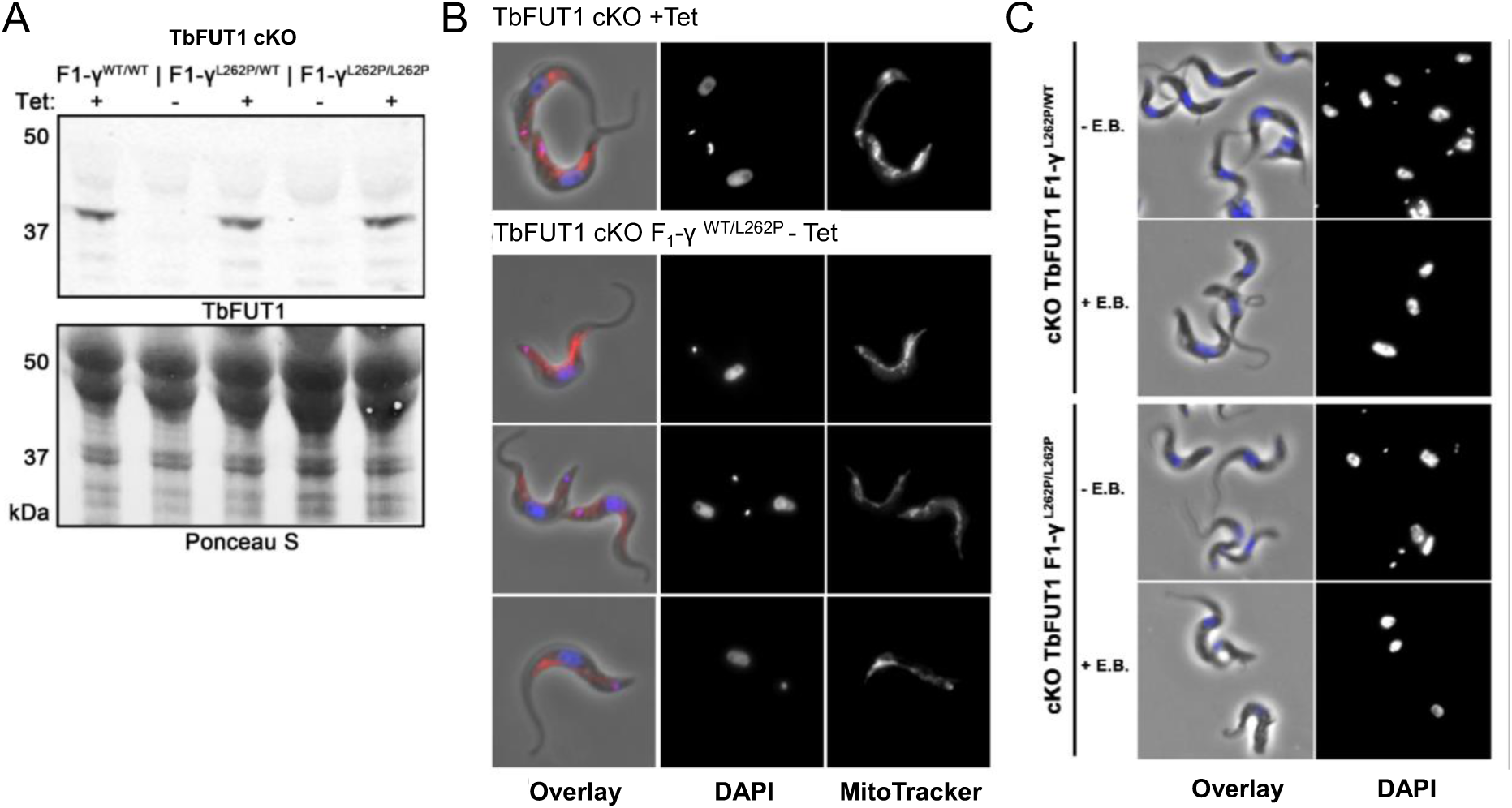
TbFUT1 is not essential for kDNA maintenance. A. Protein lysates harvested from heterozygotic (TbFUT1 cKO F_1_-γ ^WT/L262P^) and homozygous (TbFUT1 cKO F_1_-γ ^L262P/L262P^) mutants grown continuously in the presence (+ Tet) or absence (-Tet) of tetracycline were separated by denaturing PAGE gel electrophoresis, transferred to NTC membrane and analysed by Western blotting using anti-TbFUT1 (upper panel). The membrane was subjected to Ponceau S staining as a loading control (lower panel). B. Microscopy analysis of TbFUT1 cKO mutant grown in the presence of tetracycline (+ Tet) and TbFUT1 cKO F1-γ^L262P^ cells grown in the absence of tetracycline (-Tet). Cells were grown in cell culture media containing 10nM MitoTracker for 30 mins prior to fixation, and the dye is shown in red merged with DNA stained by DAPI in blue in the overlay panel C. TbFUT1 cKO F1-γ^WT/L262P^ and TbFUT1 cKO F1-γ^L262P/L262P^ cells were grown in the absence of tetracycline to ablate TbFUT1 expression and treated with (+E.B.) or without (-E.B.) 10nM ethidium bromide for 6 days. kDNA loss was assessed by microscopic analysis of fixed cells stained with DAPI.

These data show that TbFUT1 is not directly involved in kDNA maintenance. We hypothesised that dyskinetoplasty was a secondary consequence of impaired ΔΨ_m_ upon TbFUT1 depletion.

### Tandem mass tagging proteomic analysis of TbFUT1 mutants reveals reduced levels of a cohort of mitochondrial proteins

To further investigate the effects of TbFUT1 depletion, we prepared whole cell lysates of WT and TbFUT1 cKO cells grown in the presence or absence of Tet for 48 h, at which time the growth defect of the TbFUT1 cKO mutant begins to manifest (Fig. 1A) and ∼60% of cells exhibit impaired ΔΨ_m_ generation (Fig. 1C). Tandem mass tag (TMT) peptide labelling was performed in triplicate for each experimental group and the combined tryptic peptides were analysed by mass spectrometry to quantify the steady state levels of 5,738 protein groups (S1 file).

The TbFUT1 cKO mutant grown under permissive (+Tet) conditions showed ∼5-fold overexpression of TbFUT1 compared to wild-type cells (Fig. 4A, left panel). Several other protein groups also changed, such as the inhibitor of F_1_-ATPase (TbIF1) and the glucose transporter 1B (THT1) (Fig. 4A middle and right panels) and a cohort of bloodstream expression site (BES) associated proteins (Fig. S4A), presumably as adaptations to TbFUT1 overexpression. The steady state levels of F_o_-subcomplex proteins were also lower in TbFUT1 cKO cells grown +Tet relative to WT (Fig. S4B), however F_1_-subcomplex, mitochondrial ribosomal LSU and Complex I subunits remained relatively stable (Fig. S4C and D). Some of these changes will be discussed later. After 48 h under non-permissive (-Tet) conditions, TbFUT1 expression was less than in wild-type cells, but was not ablated (Fig. 4A, left panel), and some of the adaptive protein level changes to TbFUT1 overexpression did not revert to wild-type levels, e.g. THT1 and TbIF1 (Fig. 4A, middle and right panels). The comparison of wild-type and the TbFUT1 cKO mutant ±Tet proteomes is therefore complex, but was nevertheless very informative. For example, filtering the proteomes for nuclear-encoded mitochondrial proteins and non-mitochondrial proteins, as defined in (16), shows that depletion of TbFUT1 concomitantly reduced the steady state levels of many mitochondrial proteins, with TIM12 and a mitochondrial divalent cation tolerance protein as exceptions (Fig. 4B). In contrast, non-mitochondrial proteins were generally unchanged or more abundant (Fig. 4C).

**Figure 4:**
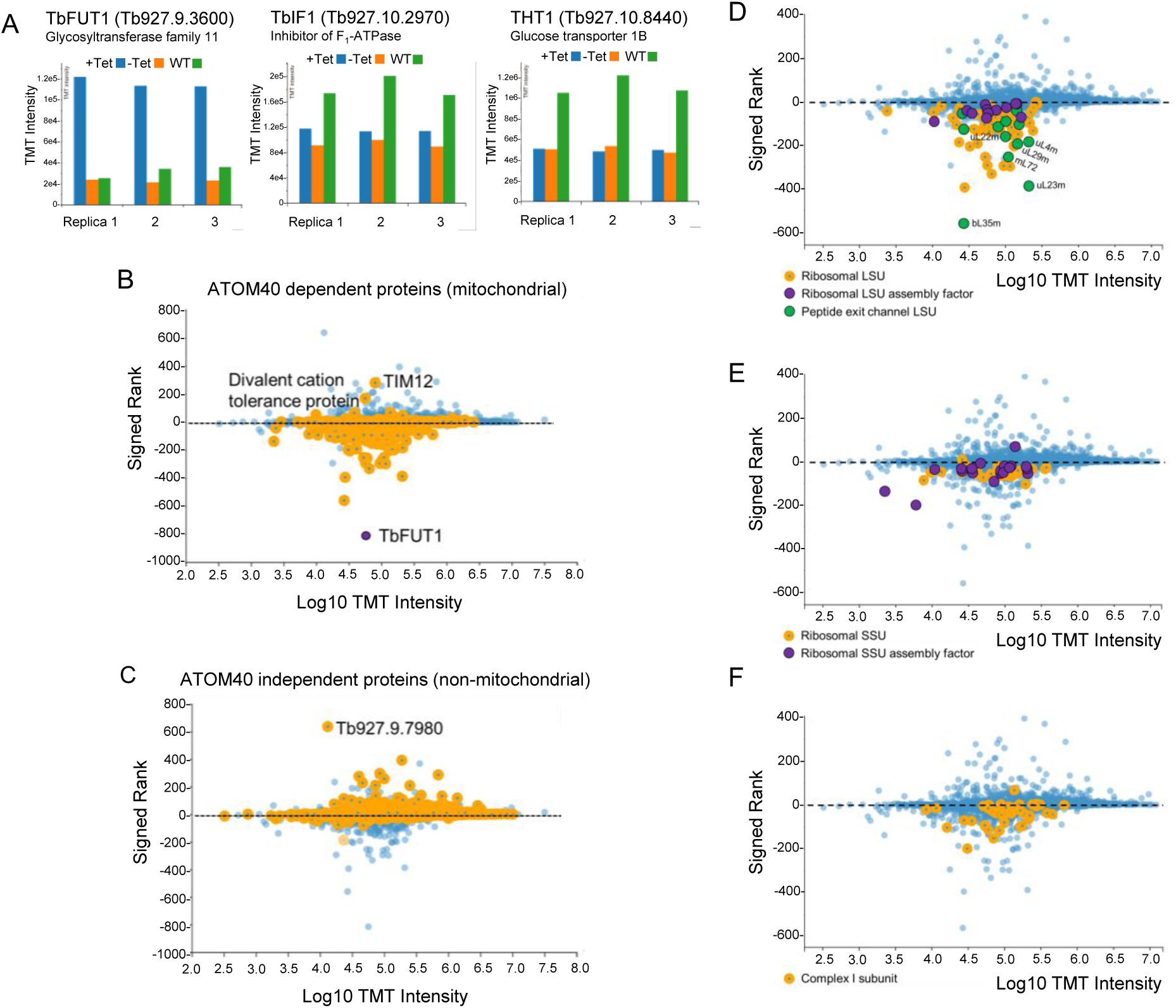
Tandem mass tagging and MS2 analysis of TbFUT1 mutants reveals reduced stead-state levels of a cohort of mitochondrial proteins. Wild-type (WT) and TbFUT1 cKO mutants were grown in the presence (+Tet) or absence (-Tet) of tetracycline for 2 days. At time of harvest cells were lysed, and extracted proteins were processed to produce reduced, alkylated tryptic peptides separately. Peptides were chemically labelled with the indicated tandem-mass tags and were combined at a 1:1 ratio following quenching of the labelling reaction. The combined peptides were fractionated by high-pH reverse phase chromatography into 80 fractions which were prepared for mass spectrometry and acquired on a Q-exactive-HF mass spectrometer. A. Steady state levels of TbFUT1 (left), TbIF1 (middle) and THT1 (right). Protein abundance is represented by TMT detection intensity. B. Steady state levels of ATOM69 dependent, mitochondrial imported proteins (orange). C. Steady state levels of non-mitochondrial imported proteins (orange) D. Steady state levels of mitochondrial ribosomal large subunit (LSU) proteins (orange) with peptide exit channel subunits indicated (green). E. Steady state levels of mitochondrial ribosomal small subunit (SSU) proteins (orange) and SSU assembly factors (purple). F. Steady state levels of mitochondrial respiratory chain Complex I subunits. Signed rank (Y-axis) calculated by comparing +Tet intensity against -Tet.

Further analysis of the TbFUT1 cKO ±Tet total proteomes revealed that the proteins most affected (reduced) by TbFUT1 depletion were components of the mitochondrial ribosomal large subunit (LSU) with minor depletions in LSU assembly factors (Fig. 4D, Table 2). Further, among these, including the most reduced proteins, are twelve LSU subunits that form the ribosomal peptide exit channel (Table 2 green text; Fig. 4D green circles) (16). Components of the ribosomal small subunit (SSU) and SSU assembly factors and respiratory chain complex I components were also reduced upon TbFUT1 depletion (Fig. 4E and F).

**Table 2.**
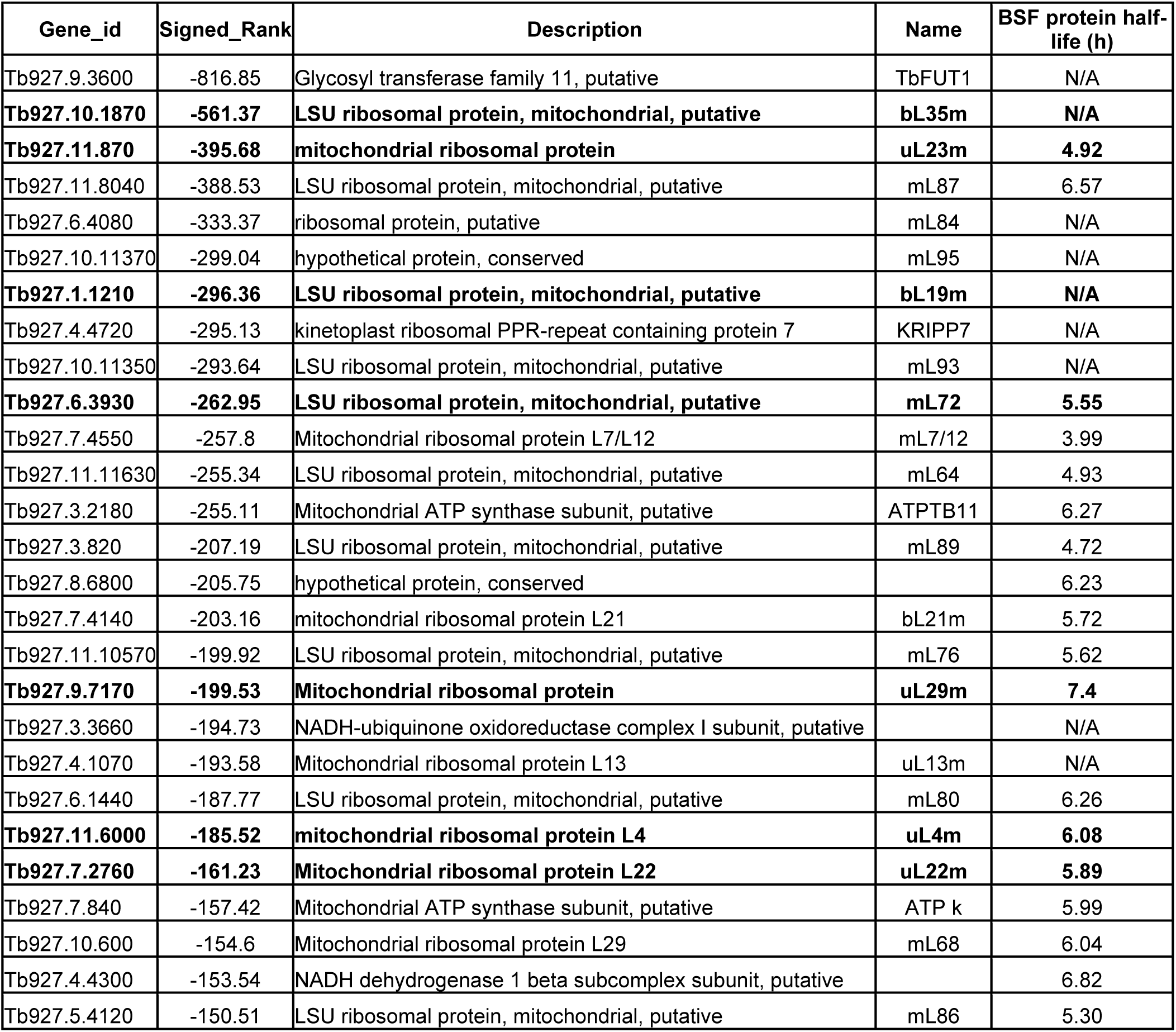
List of proteins with most reduced steady state levels following TbFUT1 KO. . TbFUT1 cKO mutants were grown in the presence (+ Tet) or absence (-Tet) of tetracycline for 2 days before TMT analysis. Signed rank calculated by comparing +Tet intensity against -Tet. Protein half-lives are indicated on the right (28). LSU components forming the peptide-exit tunnel are highlighted in bold.

Together, these results indicate that depletion of TbFUT1 correlates with a reduction in functional mitochondrial ribosome subunits, which will reduce mitochondrial protein biosynthesis and is likely to have many knock-on effects on mitochondrial and cell function.

#### Loss of TbFUT1 selectively affects F_o_F_1_-ATP synthase assembly

Since BSF *T. brucei* cells drive ΨΔm by ATP hydrolysis activity of F_o_F_1_-ATP synthase, we investigated the effect of TbFUT1 depletion on F_o_F_1_-ATP synthase levels in our TbFUT1 cKO ±Tet TMT dataset (Fig. 5A). Levels of F_o_-subcomplex and peripheral stalk subunits were reduced following TbFUT1 depletion, whereas F_1_-subcomplex proteins remained stable. To look at this in a different way, we performed native PAGE and Western blotting. Anti-Tb2 (F_o_) subunit antibodies (17) revealed similar amounts of the monomeric (M) and dimeric (D) forms of the F_o_F_1_-ATP synthase complex in wild-type cells and in the TbFUT1 cKO cells grown +Tet for 48 h and -Tet for 24 h, but with significantly reduced levels of both when grown -Tet for 48 h (Fig. 5B). Probing with anti-F_1_ β-subunit antibodies (Fig. 5B) confirmed the absence of D F_O_F_1_ 48 h after Tet withdrawal and revealed a simultaneous accumulation of unincorporated F_1_ complexes. To a lesser extent, these were also evident in the +Tet cKO cells. These data suggest that TbFUT1 is required for stable M and D F_o_F_1_-ATP synthase complex assembly and that surplus F_1_ intermediates made during TbFUT1 overexpression (+Tet) persist and cannot be incorporated into M and D F_o_F_1_-ATP synthase complexes upon TbFUT1 depletion.

**Figure 5:**
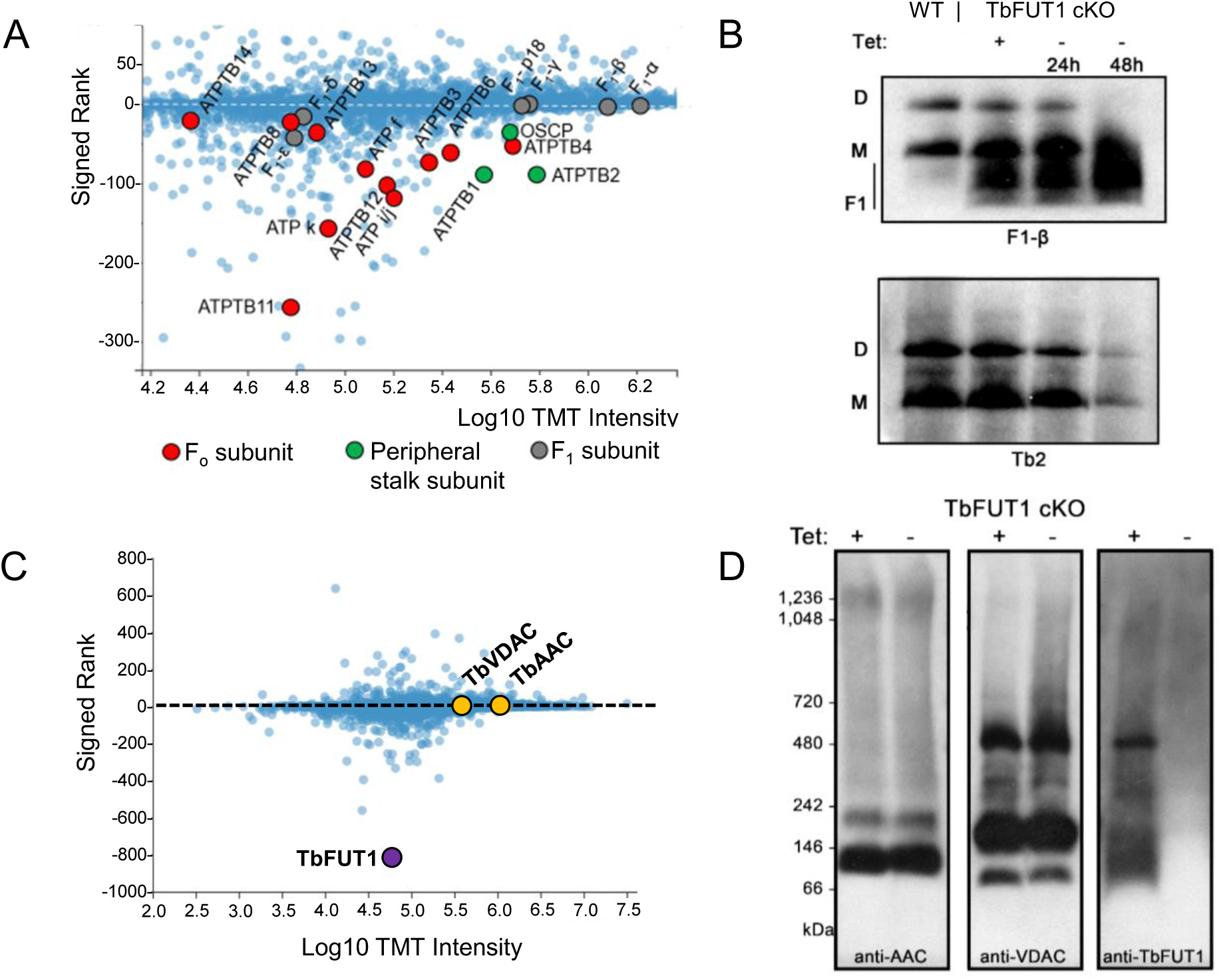
Depleting TbFUT1 inhibits the assembly of the F_O_ ATP synthase subcomplex. A. Comparative analysis of the steady state levels of F_o_-subcomplex (red), peripheral stalk (green) and F_1_-ATP (grey) synthase subunits from TbFUT1 cKO mutants grown in the presence (+Tet) or absence (-Tet) of tetracycline for 2 days. Signed rank (Y-axis) calculated by comparing +Tet intensity against -Tet. B Protein lysates harvested from wild-type (WT), TbFUT1 cKO BSF cells grown in the presence (+Tet) of Tet for 48 h, or absence (-Tet) of Tet for 24 or 48 hours were resolved by NativePAGE gel electrophoresis, transferred to a PVDF membrane, and analysed by Western blot using antibodies against F_1_-β or Tb2 proteins. The protein bands corresponding to F_1_-intermediate (F1), monomeric (M) and dimeric (D) F_o_F_1_-ATP synthase complexes are indicated on the left. C. Comparative analysis of the steady state levels of the ATP/ADP carrier protein (AAC), voltage dependent anion channel (VDAC) and TbFUT1 from TbFUT1 cKO mutants grown in the presence (+Tet) or absence (-Tet) of tetracycline for 2 days D. Proteins harvested from TbFUT1 cKO cells grown in the presence (+Tet) or absence (-Tet) of tetracycline for 3 days were resolved by NativePAGE gel electrophoresis, transferred to a PVDF membrane and analysed by Western blot using antibodies against the ATP/ADP carrier protein (AAC), voltage dependent anion channel (VDAC) and TbFUT1. A molecular weight marker is indicated on the left.

To investigate the specificity of the linkage between TbFUT1 expression and F_o_F_1_-ATP synthase complex formation, we also performed native PAGE and Western blot analysis for two unrelated mitochondrial membrane protein complexes: the mitochondrial inner membrane ADP/ATP carrier protein (AAC) and the mitochondrial outer membrane voltage dependent ion-channel (VDAC). These appear stable 48 h after TbFUT1 depletion by quantitative proteomics (Fig. 5C) and, in contrast to subunits of the F_o_F_1_-ATP synthase, the Western blot patterns for these complexes were not affected by the presence or depletion of TbFUT1 (Fig 5D). These data suggest that the import and assembly of mitochondrial membrane protein complexes, in general, are not affected by TbFUT1 and that, therefore, F_o_F_1_-ATP synthase complex assembly is specifically or selectively dependent on TbFUT1 expression.

#### Proteomic analyses of F_1_ subunit and Monomeric and Dimeric F_o_F_1_-ATP synthase complexes

To further analyse the loss of F_o_-subunits and the accumulation of F_1_-subunits in TbFUT1 overexpressing and depleted cells, respectively, proteins from wild-type and TbFUT1 cKO cells grown ± Tet for 72 h alongside the TbFUT1 cKO/F1-γ^WT/L262P^ mutant grown continuously +Tet were resolved by native PAGE in duplicate. As previously observed (Fig. 5B), wild type and TbFUT1 overexpressing (+Tet) cells contained M and D forms of the F_o_F_1_-ATP synthase complexes, as judged by anti-Tb2 (F_o_) blotting, whereas both overexpression (+Tet) or depletion (-Tet) of TbFUT1 led to the accumulation of unassembled F_1_-intermediate proteins compared to wild type, as judged by anti-F1-β blotting (Fig. 6). The duplicate native PAGE gel was fixed by Quick Coomassie and gel slices corresponding to F_1_ intermediates, monomeric and dimeric F_o_F_1_-ATP synthase complexes were excised (Fig. S5). Proteins in these slices were identified by protease digestion and LC-MS/MS analysis. The protein abundance of each F_o_F_1_-ATP synthase subunit was estimated using exponentially modified protein abundance index (emPAI) values (Table 3). F_1_-subcomplex proteins (α, β, γ, δ, ε, p18) were present in all F_1_ intermediate samples, with increased abundance in the TbFUT1 cKO ±Tet samples relative to WT. Peripheral stalk subunits (OCSP, ATPTB1 and 2) were detected in M and D complexes at similar levels in both WT and the TbFUT1 cKO +Tet samples, but were absent in the TbFUT1 cKO -Tet sample. F_o_-subcomplex proteins were present in the M and D samples of WT and TbFUT1 cKO +Tet samples (although there was quantitative variability between the samples), whereas all of the F_o_-subcomplex were notably absent in the TbFUT1 cKO -Tet sample. This included the highly hydrophobic (18, 19) mitochondrially-encoded A6 subunit which was detectable (but not quantifiable) in the M and D protein preparations of WT and TbFUT1 cKO +Tet samples, but not in the TbFUT1 cKO -Tet sample. Interestingly, the F_o_ c-ring subunit was only detected in the WT M and D extracts. As the multimeric, proton pore forming c-ring is closely associated with the A6 subunit, its absence except in WT M and D samples suggests a reduced level of A6. Further, very few F_o_-subcomplex proteins are detected in protein samples derived from the TbFUT1 depleted mutant (-Tet). Instead, only F_1_-subcomplex proteins are identified in the high molecular weight complexes in these mutants, likely deriving from multimeric F_1_-complexes associating weakly with membrane-bound proteins (17).

**Figure 6:**
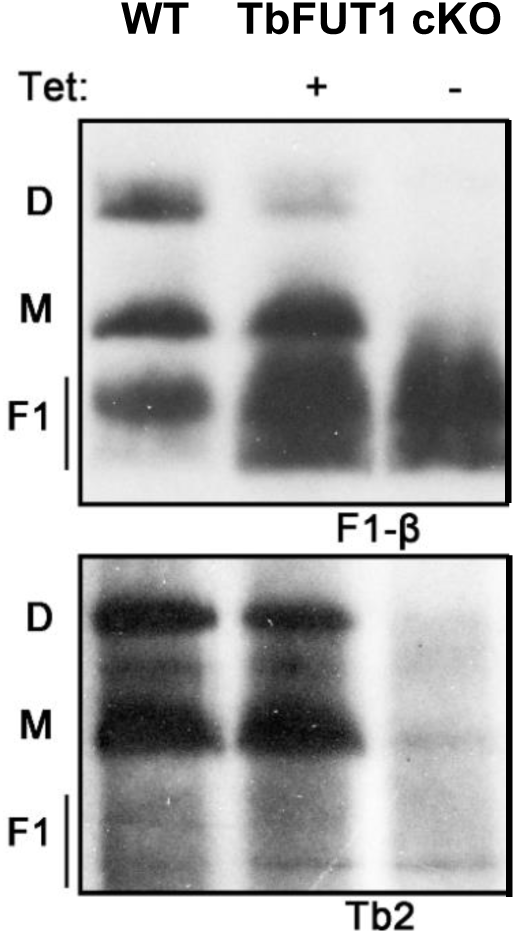
Protein identification validates the absence of F_o_-subunits from TbFUT1 cKO cells. Protein lysates harvested from wild-type (WT) and TbFUT1 cKO BSF cells grown in the presence (+T) or absence (-T) of tetracycline for 3 days were resolved by NativePAGE gel electrophoresis, transferred to a PVDF membrane and analysed by Western blot using antibodies against F_1_-β or Tb2 proteins. The protein bands corresponding to F1-intermediate (F1), monomeric (M) and dimeric (D) F_o_F_1_-ATP synthase complexes are indicated on the left and a molecular weight marker on the right. Duplicate samples were resolved, and the gel stained with Coomassie for peptide identification (Fig. S5, Table 3).

**Table 3.**
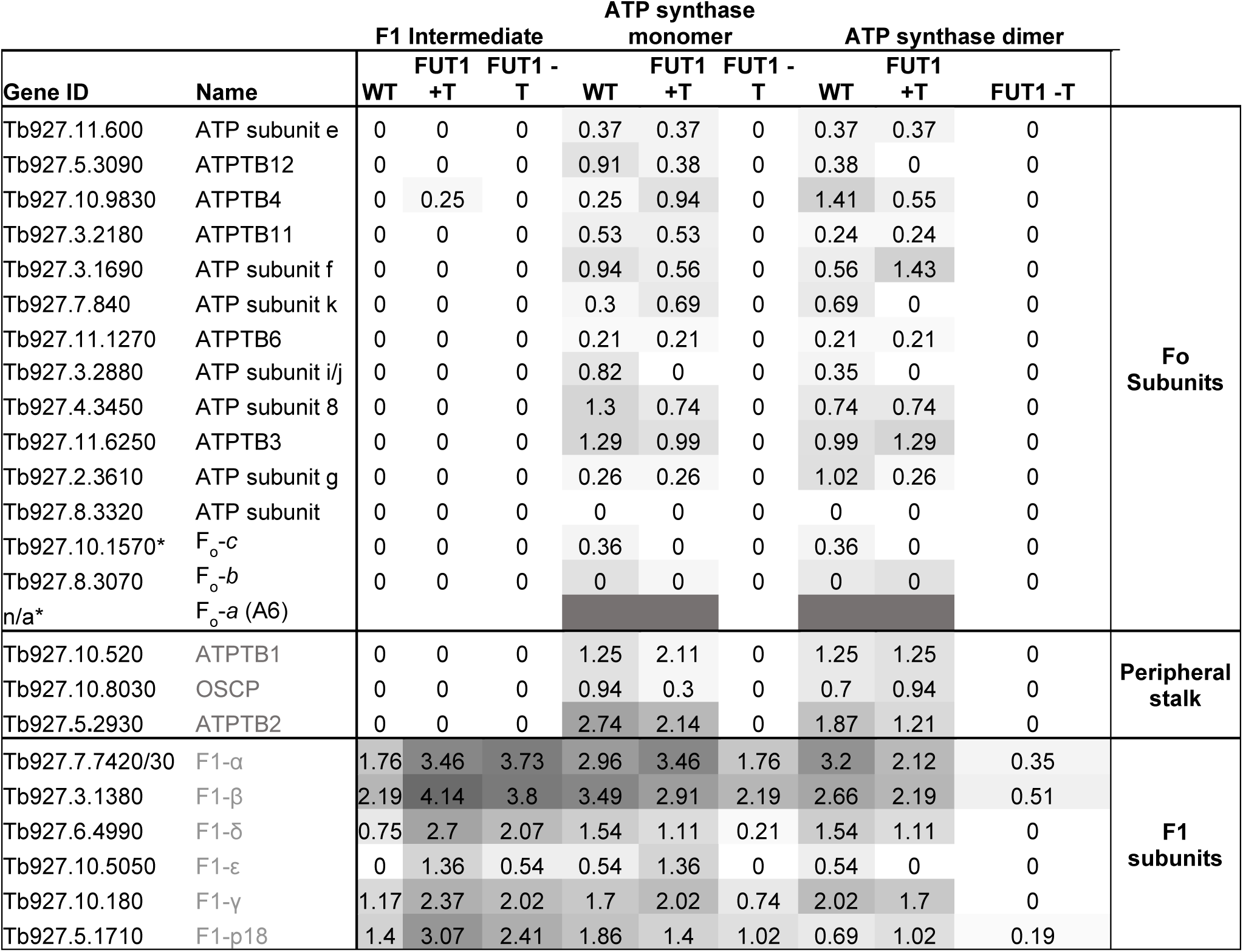
ATP synthase subunit protein identification indicates loss of F_o_ and peripheral stalk subunits by TbFUT1 KO. Protein identification of F1-intermediate, monomeric and dimeric ATP synthase complexes from wild-type (WT), TbFUT1 cKO and TbFUT1 cKO F1-γ^WT/L262P^ BSF cells grown in the presence (+Tet) or absence (-Tet) of Tet for 3 days. Proteins were resolved by NativePAGE gel electrophoresis, stained with Quick Coomassie and gel slices corresponding to the molecular mass of F_1_ intermediate (F_1_), monomeric (M) and dimeric (D) F_o_F_1_-ATP complexes. The heat map represents the EmPAI scores obtained for each of the known ATP synthase proteins represented by their TriTrypDB gene IDs. *No EmPAI scores are available for the F_o_-A6 subunit which was detected at low confidence levels. Instead, the presence of the subunit is indicated by shading in grey.

These proteomic data confirm the Western blotting data that TbFUT1 overexpression results in the accumulation of the F_1_-intermediates, that persist upon TbFUT1 depletion, and that the assembly of stable monomeric and dimeric F_o_F_1_-ATP synthase complexes is compromised upon TbFUT1 depletion.

#### All ATP synthase subunits lack detectable glycan modifications

We investigated the possibility that F_o_F_1_-ATP synthase assembly is mediated by glycosylation of one or more subunits by searching for changes to theoretical peptide masses corresponding to fucose modifications. Peptides corresponding to the F_o_F_1_-ATP synthase subunits detected in Table 3 were searched for mass increases corresponding to the addition of Fuc (dHex +146.0579), GlcNAc (HexNAc +203.079373), Galβ1-3GlcNAc (Hex(1)HexNAc(1) +365.132196) or Fucα1-2Galβ1-3GlcNAc (dHex(1)Hex(1)HexNAc(1) +511.190105). These masses were chosen based on the substrate specificity study reported in (10) which suggested that TbFUT1 prefers an acceptor glycan of Galβ1-3GlcNAc. No peptides containing any of the above modifications were detected, suggesting *T. brucei* F_o_F_1_-ATP synthase subunits are not obviously O-glycosylated with Fuc, HexNAc, Hex-HexNAc or dHex-Hex-HexNAc chains. However, the aforementioned searches are by no means exhaustive of all glycan possibilities.

#### TbFUT1 loss affects mitochondrial RNA levels

We sought to investigate the effect of TbFUT1 overexpression and depletion on mitochondrial protein synthesis, but analysis of mitochondrial protein translation in BSF cells by [^35^S]methionine radiolabelling approaches is not feasible (20). Instead, we used Real-Time quantitative PCR (RT-qPCR) analysis of a panel of mitochondrial RNA transcripts. These included pre- and post-edited transcripts for the F_o_-ATP synthase A6 subunit and the mitochondrial ribosomal SSU protein 12 (RPS12), as well as unedited 9S and 12S ribosomal RNAs and NADH-dehydrogenase (ND) subunit 1, 2 and 4 transcripts. Total RNA was extracted from wild-type cells and from the TbFUT1 cKO and TbFUT1 cKO/F1-γ^WT/L262P^ mutants grown ±Tet for 48 h. The latter replicate kDNA and generate ΔΨ_m_ independently of TbFUT1 expression (Fig. 3B).

The unedited 9S and 12S rRNA and ND subunit 1-4 transcript levels were similar to those of wild type cells in the +Tet (TbFUT1 overexpression) condition and reduced in -Tet (TbFUT1 depletion) conditions, independent of the F1-γ L262P mutation (Fig. 7A). For the pan-edited transcripts A6 and RPS12, the pre-edited transcript levels were either similar or greater than in the wild type in the +Tet (TbFUT1 overexpression) condition and the post-editing transcripts were reduced ±Tet and most dramatically for the RPS12 edited transcript under -Tet (TbFUT1 depletion) conditions (Fig. 7A and Fig. S6).

**Figure 7:**
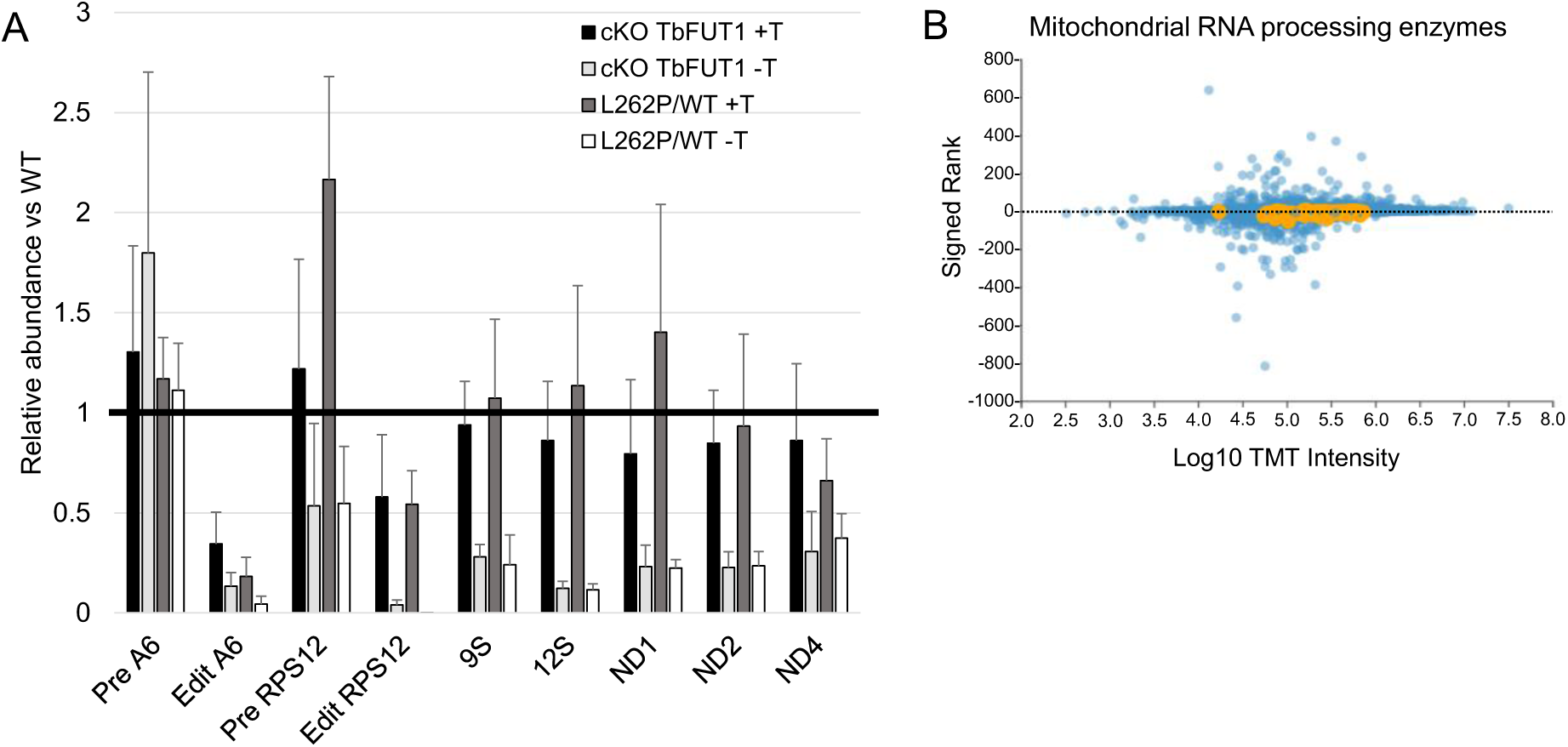
TbFUT1 cKO cells exhibit reduced mitochondrial ΔΨ_m_ impaired mitochondrial transcript editing and reduced ATP synthase assembly. A. Real-Time Quantitative PCR (RT-qPCR) analysis of RNA extracted from TbFUT1 cKO mutants grown in the presence (+) or absence (-) of tetracycline for 48 hours and TbFUT1 cKO F_1_-γ ^WT/L262P^ mutants grown continuously in the presence (+) or absence (-) of tetracycline. Nuclear encoded β-tubulin mRNA was used as internal normalisation control to generate individual ΔCt values. Relative abundance between experimental samples and a wild-type control were calculated by subtracting the ΔCt Control value from ΔCt Experimental to calculate ΔΔCt. Fold change calculated as 2^-(ΔΔ Ct). Error bars indicate the standard deviation around the means of three biological replicates, each performed in technical triplicates. B. Comparative analysis of the steady state levels mitochondrial RNA processing proteins (gene IDs obtained in (21)) from TbFUT1 cKO mutants grown in the presence (of tetracycline for 2 days. Signed rank (Y-axis) calculated by comparing +Tet intensity against -Tet.

We also searched our quantitative proteomics dataset for proteins involved in kDNA transcript editing (21). However, they were not altered (Fig. 7B), suggesting that TbFUT1 does not directly affect the assembly of mitochondrial RNA editing complexes. Together, these RT-qPCR data indicate that TbFUT1 loss reduces the levels of never edited transcripts and severely depletes levels of pan-edited A6 and RPS12 whilst overexpression of TbFUT1 particularly reduced levels of pan-edited transcripts. The effects on transcript levels following TbFUT1 depletion are not likely to be a consequence of impaired ΔΨm and kDNA replication (Fig. 1) as the F1-γ L262P mutants depleted of TbFUT1 exhibit similar levels of transcripts ± Tet (Fig. 7a, Fig. S6).

## DISCUSSION

The proposed unusual evolutionary origin (13), and the distinct GT11 clade of kinetoplastid essential mitochondrial *FUT1* gene products (10, 11), make it difficult to predict their acceptor substrates and functions. In this study we applied quantitative proteomic and biochemical approaches to better understand the effects of TbFUT1 depletion in BSF *T. brucei*.

Together, the data presented here indicate that TbFUT1 expression is required for normal mitochondrial gene transcription and translation. This view is supported by quantitative proteomics data that show reduced levels of mitochondrial protein complexes containing kDNA-encoded subunits in TbFUT1 depleted cells (Fig. 4 and 5, Table 2). The disruption of normal mitochondrial gene transcription and translation upon TbFUT1 depletion has multiple effects. We propose these are manifested by the failure to make (A6 subunit) or stabilise F_o_-subunits, the consequent failure to assemble mitochondrial monomeric and dimeric F_o_F_1_-ATP synthase complexes (but not other mitochondrial membrane protein complexes lacking kDNA-encoded products), the corresponding collapse of mitochondrial ΔΨ_m_ potential and, ultimately, dyskinetoplasty and cell death. The reduced mitochondrional ribosome subunit steady state levels and rRNA levels following TbFUT1 depletion further imply that TbFUT1 contributes to mitochondrial gene transcription and translation. The F_o_-independent mode of ΔΨ_m_ generation in cells containing the F_1_-γ ^L262P^ mutation explains why this mutation rescues TbFUT1 depletion; i.e., mitochondrial gene expression and translation are no longer necessary in these cells. Finally, the retention of kDNA following TbFUT1 loss in cells containing F_1_-γ ^L262P^ indicates that TbFUT1 does not directly mediate mitochondrial DNA synthesis/replication.

While we have added to our understanding of how TbFUT1 depletion leads to cell death in BSF *T. brucei,* the mechanism(s) by which TbFUT1 maintains mitochondrial transcription and/or translation are still unknown. The process of mitochondrial gene expression and translation is complex, and involves multiple RNA binding proteins, editing complexes and guide RNAs (21), and is further interconnected with kDNA replication (22) and mitochondrial translation (23). Therefore, precisely how mitochondrial fucosylation exerts global changes in mRNA levels is unclear, particularly as RNA processing and editing enzyme steady states were unchanged (21) (Fig. 6B), suggesting they are still forming functional complexes.

Our search for endogenous TbFUT1 substrates continues, and further clues may emerge from identifying TbFUT1 binding partners. Existence of the latter are suggested from the native PAGE anti-TbFUT1 western blot in (Fig. 5D). Thus, overexpression and/or depletion of TbFUT1 may affect the activities of partner proteins with possible roles in RNA processing. Alternatively, we cannot exclude the possibility that mitochondrial RNA levels and/or editing are directly regulated by fucosylation. The glycosylation of RNA has been described in mammalian cells and includes fucose modifications (24). In that case, the synthesis of “glycoRNas” requires canonical biosynthetic enzymes contributing to *N*-glycosylation through the secretory pathway and not a mitochondrial process. Therefore, the mechanism by which TbFUT1 regulates mitochondrial transcript levels requires further investigation.

TbFUT1 is normally detected at low abundance in BSF and PCF *T. brucei* (25). Our data indicate that the >5 fold overexpression of TbFUT1 in our conditional null mutants under permissive conditions (Fig. 4A, left panel) perturbs mitochondrial function, albeit more subtly than TbFUT1 depletion. Relative to WT, our TbFUT1 overexpressing cells have reduced pan-edited of the *A6* transcript (Fig. 7A), more F_1_-intermediate subcomplexes (Fig. 5B, Fig. 6A, Table 3) and reduced levels of F_o_-ATP synthase and peripheral stalk subunits (Fig. S4B). Our TbFUT1 overexpressing mutants generate a sufficient ΔΨ_m_ (Fig. 1C) via F_o_F_1_-ATP synthase to permit normal cell growth. Yet the reduced assembly of F_o_F_1_-ATP synthase in TbFUT1 overexpressing mutants may exert changes to other, associated proteins. For example, the inhibitor of F_1_-ATPase (TbIF1) that binds to the F_1_-ATP moiety to inhibit ATP hydrolysis (26) is reduced to ∼50% WT levels in TbFUT1 cKO mutants ± Tet (Fig. 4A). The reduction of TbIF1 in TbFUT1 cKO mutants may be linked to the reduced levels of fully assembled ATP synthase. Additionally, the levels of the main glucose transporter 1B (THT1) of BSF *T. brucei* (27) (Fig. 4A, right panel) are similarly reduced to ∼50% of wild-type levels in TbFUT1 cKO mutants ± Tet. BSF *T. brucei* synthesise the majority of their energy through glucose uptake and glycolysis, importing ATP into the mitochondrial matrix via TbAAC (Fig. 5C). Unlike TbIF1 however, THT1 this is not directly associated with the ATP synthase and its modulated abundance may instead be linked with impaired ATPase activity, as a mechanism to limit ATP production from glycolysis.

We discovered through our proteomics survey that the most depleted proteins, after TbFUT1 itself, under non-permissive conditions were of the mitochondrial ribosomal LSU (Fig. 4D, Table 2). This implies reduced mitochondrial protein synthetic capacity and reduced stabilisation of mitochondrial complexes containing mitochondrially-encoded subunits, such as F_o_-ATP synthase (Fig. 5A and B). Curiously, unlike the steady state levels of F_o_-ATP synthase proteins, the mitochondrial ribosome subunits were not altered by >5 fold TbFUT1 overexpression (Fig. S4C). An obvious explanation for mitochondrial ribosome destabilisation is the loss of kDNA and thus reduced levels of kDNA transcripts encoding key mitochondrial ribosome factors e.g. pan-edited RPS12 or 9S/12S rRNAs, following TbFUT1 depletion (Fig. 7A). However, the similar depletion of pan-edited and never-edited transcripts in TbFUT1 cKO and TbFUT1 cKO F1-γ^WT/L262P^ mutants occurs at similar rates, despite the latter retaining kDNA (Fig. S6). Further, the general loss of kDNA encoded factors does not correlate with the more pronounced depletion of LSU over SSU subunits, despite their similar rates of protein turnover (28). Therefore, LSU destabilisation may arise by another, undefined mechanism.

An alternative possibility for the depletion of LSU proteins in particular may stem from the fact that Trypanosomal mitoribosomes are highly proteinaceous and relatively lacking in rRNA (16). Ribosomal protein shells form early during the maturation process, assembling almost independently of the rRNA core (29). The 12S rRNA elements that form key functional regions of the mitoribosomal LSU, including the peptide exit channel and peptidyl transfer channel, are held in immature conformations stabilized by mitochondrial LSU assembly factors (mt-LAFs), prior to 12S rRNA insertion (30). Therefore, LSU depletion could be because of impaired mt-LAF activity. Yet a survey of mt-LAFs reveals more minor reductions in protein abundance following TbFUT1 loss (Fig. 3D and E), relative to LSU components. Interestingly, RNAi of such mt-LAFs impairs both LSU assembly but also results in a corresponded 60% decrease in both 9S and 12S rRNA (30). The loss of 9S and 12S rRNA in our TbFUT1 mutants may therefore be compounded by reduced LSU stability, contributing to rRNA loss and impaired translation. This may be an alternative explanation for the TbFUT1 dependence of RNA levels in TbFUT1 cKO F1-γ^WT/L262P^ mutants, other than the direct modification of transcripts with fucose.

Interestingly, LSU subunits bL35m (Tb927.10.1870) and uL23m (Tb927.11.870), along with ten other proteins form the peptide exit tunnel region of the LSU (16) were amongst the most depleted by TbFUT1 loss (Fig 4D, Table 2). Why the exit tunnel region is particularly affected by TbFUT1 depletion is unclear. The cause may be by the ∼40% loss of mt-LAF2, which aids in the assembly of the peptide exit tunnel stabilising the region later occupied by 12S rRNA. An alternative explanation may be the position of the exit tunnel, a region tightly associated with the inner mitochondrial membrane for efficient protein delivery. Co-translational insertion of F_o_-A6 is mediated by Oxa1 in yeast (31) and human OXA1L (32), suggesting a similar role in *T. brucei*. Two Oxa1-like proteins are encoded by *T. brucei* (33), where one is expressed in BSF and PCF life cycle stages (Tb927.11.6150) and the other restricted to PCF cells (Tb927.9.10050) (25); their function is yet to be investigated experimentally. Unfortunately, neither protein was detected in our proteomics analysis, and whether FUT1 depletion affected their abundance therefore remains unknown.

In summary, we demonstrate that TbFUT1 has a critical role in selective mitochondrial complex assembly by regulating the expression of kDNA encoded transcripts and/or proteins. Depletion of TbFUT1 impairs mitochondrial membrane potential via loss of the F_o_-ATP synthase subcomplex, whilst the reduced levels of mitochondrial rRNA and proteins likely impair mitochondrial protein translation capacity further. Further investigation is necessary to discover the substrate for mitochondrial fucosylation and uncover the precise mechanism by which mitochondrial complex assembly is regulated by TbFUT1.

## Materials and Methods

### Cultivation of Trypanosomes—Trypanosoma brucei brucei

Lister strain 427 bloodstream form parasites, expressing VSG variant 221 (MiTat1.2) and transformed to stably express T7 polymerase and the tetracycline repressor protein under G418 antibiotic selection, were used in this study. This genetic background will be referred to from hereon as wild-type (WT). Cells were cultivated in HMI-11 medium containing 2.5 μg/mL G418 at 37 °C in a 5% CO_2_ incubator as described in (34).

### Growth curves

Growth of parasites lacking TbFUT1(10) was analysed by determining cell counts for BSF TbFUT1 cKO. Cells were washed in media without tetracycline (tet) and inoculated at a concentration of 5 × 10^3^ parasites/ml. Parasites were grown plus and minus tetracycline for 72 h and counted every 24 h using a Neubauer chamber in a phase contrast microscope.

### Immunofluorescence microscopy

Late log phase *T. brucei* bloodstream form cells were fixed in 4% PFA/PBS in solution at a concentration of 5 × 10^6^ cells/ml. When using MitoTracker Red CMX Ros, cell cultures were spiked with a 25 nM concentration over 20 min before harvesting. Fixed cells were mounted using ProLong^TM^ Gold mountant with DAPI (Thermo Fisher) prior to microscopy to stain nuclear and kinetoplast DNA.

### ΔΨm measurement

The ΔΨm of wild-type and TbFUT1 cKO cells grown in permissive and non-permissive conditions was estimated using the cell permeant red-fluorescent dye tetramethyl rhodamine (TMRE, Thermo Fisher Scientific T669). Fluorescence intensity is proportionally dependent on the ΔΨm values. Equal number of cells (5 × 10^6^) were harvested for each time point and resuspended in the culture medium containing 60-nM TMRE for 30 min at 37 °C. Cells were centrifuged at 1,000 g for 10 min at room temperature, resuspended in 1 ml of 1× PBS (see composition above) and immediately analysed by flow cytometry using a CytoFlex Flow Cytometer instrument and its blue laser (488 nm) with the band pass PE filter (585/15). For each sample, 10,000 events were collected. Data were evaluated using Cytoflex software (Beckman). The TMRE signal corresponding to TbFUT1 cKO cells was normalized to that of TMRE labelled wild type and unstained controls, and expressed in percentage.

### DNA Isolation and Manipulation

Plasmid DNA was purified from *Escherichia coli* DH5α competent cells (MRC PPU Reagents & Services, Dundee) using a Qiagen Miniprep kit. Gel extraction and reaction clean-up was performed using Qiaquick kits (Qiagen). Custom oligonucleotides were obtained from Thermo Fisher. *T. brucei* genomic DNA was isolated from ∼5 × 10^7^ BSF cells using lysis buffer containing 100 mM Tris-HCl (pH 8.0), 100 mM NaCl, 25 mM EDTA, 0.5% SDS, and 0.1 mg/mL proteinase K (Sigma) by standard methods.

### Transformation of bloodstream form T. brucei

Gene replacement constructs designed to replace a single allele of subunit F_1_-γ by homologous recombination to confer F_1_-γ^L262P^ point mutation were a kind gift from Prof. Schanufer(7). Each construct containing either puromycin acetyltransferase (*PAC*) or blasticidin deaminase (*BSD*) drug resistance cassettes were digested with appropriate restriction enzymes to linearize, precipitated, washed with 70% ethanol, and re-dissolved in sterile water. The linearized DNA was electroporated into *T. brucei* bloodstream form cells (Lister strain 427, variant 221) that were stably transformed to express T7 RNA polymerase and the tetracycline repressor protein under G418 selection. Cell transformation was carried out as described previously (34–36). The heterozygotic mutant (TbFUT1 cKO F_1_-γ^L262P/WT^) was generated by electroporation of the TbFUT1 cKO mutant in the presence of linear *PAC* cassette in the absence of tetracycline. A puromycin resistant clone containing a single L262P mutant allele was ultimately achieved by removing tetracycline from the culture medium at the time of transfection. The F1-γL262P double allele mutant was then readily generated by transforming the TbFUT1 cKO/F1-γWT/L262P cell line with an L262P mutagenesis cassette conferring blasticidin resistance Second round electroporation was performed to replace the remaining allele with the *BSD* cassette, generating a homozygous mutant (TbFUT1 cKO F_1_-γ^L262P/L262P^). Allelic replacement of endogenous subunit *γ* was confirmed by DNA sequencing using oligonucleotide primers SMD337/8. The cell lines were tested for kDNA independence by treating with 10 nM ethidium bromide for two passages, resulting in the generation of viable akinetoplastic cells.

### Southern blotting

10 μg aliquots of genomic DNA isolated from TbFUT1 cKO mutants at 24, 48 and 72 h following removal of tetracycline from the cell culture media were digested with EcoRI, resolved on a 0.8% agarose gel and transferred onto a Hybond-N positively charged membrane (GE Healthcare, UK). Oligonucleotide primers were used to amplify a Maxi-probe (SMD359/6) and Hexose transporter control-probe (SMD361/2) from genomic DNA template. Highly sensitive DNAprobes labelled with digoxigenin-dUTP were generated using the PCR digoxigenin probe synthesis kit (Roche Applied Science) according to the manufacturer’s recommendations. The control-probe was first hybridized overnight at 42 °C followed by washing of the membrane and a second overnight hybridisation with the Maxi-probe. Detection was performed using alkaline phosphatase-conjugated anti-digoxigenin Fab fragments and the chemiluminescent substrate CSPD (Roche Applied Science). ImageJ analysis was performed to calculate the relative intensity of Maxi-probe detected bands with the corresponding control-probe detected bands.

### RNA isolation and RT-qPCR analysis

To assess the expression of kinetoplast encoded transcripts, wild-type (WT), TbFUT1 cKO and TbFUT1 cKO F_1_-γ^L262P/WT^ cells grown under permissive and non-permissive conditions and RNA was extracted from 1 ×10^7^ cells using RNeasy RNA extraction kit (Qiagen). RT-qPCRs were performed using Luna® Universal qPCR Master Mix (NEB) and acquired on a QuantStudio 3 system (Applied Biosystems) according to manufacturer’s instructions. Oligonucleotide primer pairs amplifying < 150 nucleotide products of pre-edited A6 (SMD392/3) and RPS12 (SMD406/7), pan-edited A6 (SMD394/5) and RPS12 (SMD408/9), ND1 (SMD432/3), ND2 (SMD434/5), ND4 (SMD422/3), 9S rRNA (SMD426/7) and 12S rRNA (SMD428/9) were used (20). A housekeeping β-tubulin mRNA control (SMD398/9) was included as a normalisation control to generate individual ΔCt values. Relative abundance between experimental samples and the wild-type control were calculated by subtracting the ΔCt Control value from ΔCt experimental to calculate the ΔΔCt value. Relative abundance as compared to wild-type levels is calculated as 2^-(ΔΔCt).

### SDS-PAGE and NativePAGE Western blotting

To confirm the tetracycline inducible expression of TbFUT1, cells were grown in the presence or absence of tetracycline. 1 x 10^7^ cells were lysed in 25mM Tris, pH 7.5, 100 mM NaCl, 1% Triton X-100 and solubilised in 1xSDS sample buffer containing 0.1 M DTT by heating at 55°C for 20 min. Aliquots corresponding to 5×10^6^ cells per sample, were subjected to SDS-PAGE on NuPAGE bis-Tris 10% acrylamide gels (Invitrogen) and transferred to a nitrocellulose membrane (Invitrogen). Ponceau staining confirmed equal loading and transfer. The blot was further probed with anti-TbFUT1 rabbit polyclonal antibody (10) in a 1:1,000 dilution. Detection was carried out using IRDye 800CW conjugated goat anti-rat IgG antibody (1:15,000) and LI-COR Odyssey infrared imaging system (LICOR Biosciences, Lincoln, NE). To investigate mitochondrial protein complex formation, aliquots of 2 × 10^7^ cells were harvested, washed with 1 mL 1x phosphate buffered saline (PBS) + 6 mM glucose, resuspended in 125 uL of SoTE buffer (0.6 M sorbitol, 25 mM Tris-HCl, pH7.5, 2 mM Ethylenediaminetetraacetic acid (EDTA) and 125 uL of SoTE + 0.03% Digitonin (SoTE-D) added and mixed by inverting once. The suspension was centrifuged at 4,500 g at 4°C for 3 min and the pellet containing the mitochondrial enriched fraction was resuspended in 50 uL of lysis buffer of 50 mM Tris-HCl, pH 7.4, 150 mM NaCl containing 0.5% digitonin and incubated on ice 20 min. After centrifugation at 16,000 g, 4°C for 20 min, the supernatant was removed. Cell equivalent to 1 × 10^7^ cells, were subjected to NativePAGE (Invitrogen) and transferred to a PVDF membrane (Invitrogen) followed by immunoblotting using anti-TbFUT1(10), anti-TbVDAC, anti-TbAAC antibody, anti-F_1_β and anti-Tb2 antibodies (37) (all antibodies were generously donated by Dr. Zikova) (Chromotek, 9E1) diluted 1:1,000. The blot was then developed by ECL using an HRP-conjugated secondary antibody (Sigma, A9037, 1:3,000).

### Mass spectrometry protein identification

To identify the protein composition of bands corresponding to the F_1_-intermediate, monomeric and dimer F_o_F_1_-ATP synthase, 1 × 10^7^ cells of wild type and TbFUT1 cKO mutants grown in permissive or non-permissive conditions for 72 h were harvested, mitochondrial fraction enriched and resolved by NativePAGE analysis. Whole lanes containing wild type and TbFUT1 cKO mutant samples were cut identically into 3 slices corresponding to the molecular weight of the complexes of interest. Gel slices were submitted to protein ID and the gel pieces were dried in Speed-vac (Thermo Scientific) for in-gel reduction with 0.01 M dithiothreitol and alkylation with 0.05 M iodoacetamide (Sigma) for 30 min in the dark. The gel slices were washed in 0.1 M NH_4_HCO_3_, and digested with 12.5 μg/mL modified sequence grade trypsin (Roche) in 0.02 M NH4HCO3 for 16 h at 30°C followed by Asp-N digestion to facilitate processing and detection of the highly hydrophobic A6 subunit (18, 19). Samples were dried and re-suspended in 50 μL 1% formic acid and then subjected to liquid chromatography on Ultimate 3000 RSLC nano-system (Thermo Scientific) fitted with a 3 Acclaim PepMap 100 (C18, 100 μM × 2 cm) and then separated on an Easy-Spray PepMap RSLC C18 column (75 μM × 50 cm) (Thermo Scientific). Samples (15μL) were loaded in 0.1% formic acid (buffer A) and separated using a binary gradient consisting of buffer A and buffer B (80% acetonitrile, 0.1% formic acid). Peptides were eluted with a linear gradient from 2 to 35% buffer B over 70 min. The HPLC system was coupled to a Q-Exactive Plus Mass Spectrometer (Thermo Scientific) equipped with an Easy-Spray source with temperature set at 50°C and a source voltage of 2.0 kV.

### Serial block fake scanning electron microscopy

BSF TbFUT1 cKO cells were grown in the presence or absence of tetracycline for 77h and fixed in 2.5% glutaraldehyde. Following fixation, samples were post-fixed in 1% (w/v) osmium tetroxide in PBS for 30 min at room temperature and en bloc stained with 1% (w/v) aqueous uranyl acetate. Following dehydration through an acetone series, samples were embedded in epoxy resin. Ultrathin (70-nm) sections were post-stained with 2% uranyl acetate and lead citrate and imaged on a Tecnai G2 transmission electron microscope (FEI, Hillsboro, OR, USA). Resin blocks were also imaged on a Zeiss Gemini/Merlin scanning electron microscope, and serial sections were generated using a Gatan 3view serial sectioning system. Image resolution was 8000 × 8000 pixels with a pixel size of 3 nm2 and a z-resolution of 100 nm. Initial data processing was performed using IMOD. Nine cells in S phase of the cell cycle were selected from TbFUT1 cKO – Tet samples for kDNA analysis. Each cell was then segmented and reconstructed using Amira software (versions 5.3.3 to 6.1, FEI, Eindhoven, Netherlands). Individual mitochondrial volumes were calculated for eight 2K1N cells from TbFUT1 cKO mutants grown in the presence or absence of tetracycline.

### Tandem mass tagging labelling and quantitative proteomics analysis

Wild-type and TbFUT1 cKO cells grown in permissive and non-permissive conditions for 48 h and 4 ×10^7^ cells each were harvested by centrifugation. Cells were washed twice in Trypanosoma dilution buffer (TDB) and the final washed pellets were resuspended in 25 uL 0.1 M TEAB, and a further 25 uL of 0.1 M TEAB + 10% SDS added and mixed to homogenise samples.

### S-Trap processing of samples

Samples (equivalent of 250 µg) were processed using S-trap mini protocol (Protifi) as recommended by the manufacturer with little modification. After, application of the samples on the S-trap mini spin column, trapped proteins were washed 5 times with S-TRAP binding buffer. A double digestion with trypsin (1:40) was carried out first overnight at 37°C in 160 µl of TEAB at a final concentration of 50 mM, and then for another 6 hrs (1:40) in 110 µl TEAB. Elution of peptides from M S-trap mini spin column was achieved by centrifugation at 1000 x g for 1 min by adding 160 µl of 50 mM TEAB, then 160 µl of 2% aqueous formic acid and finally 160 µl of 50% acetonitrile/0.2% formic acid. Resulting tryptic peptides were pooled, dried a couple of times, to reduce acidity, resuspended in 50 mM TEAB and quantified using Pierce Quantitative fluorometric Peptide Assay (Thermo Scientific).

### TMT Labelling and high pH reverse phase fractionation

Tryptic peptides (20 µg, each sample) were dissolved in 100 µl of 100 mM TEAB. TMT labelling was performed according to the manufacturer’s instructions (ThermoFisher Scientific). The different TMT-6 plex labels (Thermo Fisher Scientific) were dissolved in 41µL of anhydrous acetonitrile, and each label is added to a different sample. The mixture was incubated for 1 hour at room temperature and labelling reaction was stopped by adding 8µl of 5% hydroxylamine per sample. Following labelling with TMT, samples were checked for labelling efficiency, then mixed, desalted, and dried in a speed-vac at 30°C. Samples were re-dissolved in 200 µl ammonium formate (10 mM, pH 9.5) and peptides were fractionated using High pH RP Chromatography. A C18 Column from Waters (XBridge peptide BEH, 130Å, 3.5 µm 2.1 × 150 mm, Waters, Ireland) with a guard column (XBridge, C18, 3.5 µm, 2.1×10mm, Waters) were used on an Ultimate 3000 HPLC (Thermo-Scientific). Buffers A and B used for fractionation consist, respectively, of (A) 10 mM ammonium formate in milliQ water pH 9.5 and (B) 10 mM ammonium formate, pH 9.5 in 90% acetonitrile. Fractions were collected using a WPS-3000FC auto-sampler (Thermo-Scientific) at 1minute intervals. Column and guard column were equilibrated with 2% Buffer B for twenty minutes at a constant flow rate of 0.2ml/min. 190 µl of TMT labelled peptides were injected onto the column, and the separation gradient was started 1 minute after the sample was loaded onto the column. Peptides were eluted from the column with a gradient of 2% Buffer B to 20% Buffer B in 6 minutes, then from 20% Buffer B to 45% Buffer B in 51 minutes, finally from 45% buffer B to 100% Buffer B within 1 min. The Column was washed for 15 minutes in 100% Buffer B to remove hydrophobic peptides. The fraction collection started 1 minute after injection and stopped after 80 minutes (total 80 fractions, 200µl each). To acidify the eluting peptides, 30 µl of 10% formic acid was added to each of the 80 fractionation vials. Total number of fractions concatenated was set to 20.

### LC-MS Analysis

Analysis of peptides was performed on a Q-exactive-HF (Thermo Scientific) mass spectrometer coupled with a Dionex Ultimate 3000 RS (Thermo Scientific). LC buffers were the following: buffer A (0.1% formic acid in Milli-Q water (v/v)) and buffer B (80% acetonitrile and 0.1% formic acid in Milli-Q water (v/v). Peptides from each fraction were resuspended in 50 µl 1% formic acid and aliquots of 4 μL were loaded at 10 μL/min onto a trap column (100 μm × 2 cm, PepMap nanoViper C18 column, 5 μm, 100 Å, Thermo Scientific) equilibrated in 0.1% TFA. The trap column was washed for 6 min at the same flow rate with 0.1% TFA and then switched in-line with a Thermo Scientific, resolving C18 column (75 μm × 50 cm, PepMap RSLC C18 column, 2 μm, 100 Å) equilibrated in 5% buffer B for 17 min. The peptides were eluted from the column at a constant flow rate of 300 nl/min with a linear gradient from 5% buffer B (for Fractions 1-10, 7% for Fractions 11-20) to 35% buffer B in 125 min, and then to 98% buffer B in 2 min. The column was then washed with 98% buffer B for 20 min and re-equilibrated in 5% or 7% buffer B for 17 min. The column was maintained at a constant temperature of 50°C. Q-exactive HF was operated in data dependent positive ionisation mode. The source voltage was set to 2.7 Kv and the capillary temperature was at 250°C. A scan cycle comprised MS1 scan (m/z range from 335-1600, with a maximum ion injection time of 50 ms, a resolution of 120 000 and automatic gain control (AGC) value of 3×10^6^) followed by 15 sequential dependant MS2 scans (resolution 60000) of the most intense ions fulfilling predefined selection criteria (AGC 1×10^5^, maximum ion injection time 200 ms, isolation window of 0.7 m/z, fixed first mass of 100 m/z, exclusion of unassigned, singly and >6 charged precursors, peptide match preferred, exclude isotopes on, dynamic exclusion time of 45 s). The HCD collision energy was set to 32% of the normalized collision energy. Mass accuracy is checked before the start of samples analysis. Spectra of MS1 and MS2 scans were collected respectively in profile and centroid mode.

### TMT data analysis

MS/MS data were analysed for protein identifications using MaxQuant 1.6.14 (38) with the in-built Andromeda search engine. The raw files were searched against the *T. brucei* TREU927 proteome downloaded from TriTrypDB (39) supplemented with the BES1/TAR40 protein sequences downloaded from NCBI (accession number: FM162566). The mass tolerance was set to 4.5 ppm for precursor ions and trypsin set as the proteolytic enzyme with two missed cleavages permitted. Carbamidomethyl on cysteine, was set as fixed modifications. Oxidation of methionine and Acetylation of Protein N-term were set as variable modifications. The false-discovery rate for protein and peptide level identifications was set at 1%, using a target-decoy based strategy. The minimum peptide length was set to seven amino acids. “Reverse Hits”, “Only identified by site” and “Potential contaminant” identifications were filtered out. Only protein groups with at least two unique peptide sequences and Andromeda protein score greater than 1 were selected for further quantification. For the differential protein expression analysis, the TMT values were analysed with the ProtRank (40) package by comparing the three wild-type control samples versus the TbFUT1 cKO cells grown in permissive and non-permissive conditions. Briefly, ProtRank analyses the comparisons that involve a missing value separately from those that do not involve a missing value. The logarithmic fold changes and their magnitude relative to other protein fold changes are computed for comparisons without missing values, all comparisons where a zero value changes in a positive value are assigned the same relatively high virtual rank, and all comparisons where a positive value changes in a zero value are assigned the same relatively low rank. The analysis outputs were visualised using a web app available at https://sam-d-p.pages.dev for the control vs permissive comparison, available at https://sam-p-m.pages.dev for the permissive vs non-permissive comparison and available at https://sam-d-m.pages.dev for the control vs non-permissive comparison. We also visualized the signed-rank and FDR of the: 1) control vs permissive and 2) permissive vs non-permissive conditions available at https://sam-all.pages.dev. The web applications were developed using a web template described in (25).

The analysis pipeline is available in GitHub (to be defined) and deposited in Zenodo (to be defined). The analysis pipeline was implemented in python using the SciPy packages (https://www.scipy.org/) (41) and Jupyter notebook (https://jupyter.org/).

### Data availability

The mass spectrometry proteomics data have been deposited to the ProteomeXchange Consortium via the PRIDE partner repository with the dataset identifier (to be defined) for the TMT experiment and identifier (to be defined) for the gel band identifications.

## Acknowledgements

We thank Achim Schnaufer for kindly providing the L262P mutagenesis plasmids and providing invaluable experimental advice and expert review of the manuscript, Alena Zikova for kindly providing antibodies against AAC, VDAC, F_1_-β and Tb2 and providing helpful advice, Marta Garcia Sanchez for assisting with live cell flow cytometry and Giulia Bandini, Sebastian Damerow and Lucia Güther for the generation and skilled investigation of TbFUT1 mutants.

