## Supplemental Files for "Association of mitochondrial fucosyltransferase TbFUT1 with the assembly of the mitochondrial F_o_F_1_-ATP synthase in bloodstream form *Trypanosoma brucei*"

*Supporting information*

S1 File **List of proteins identified in TMT analysis in 1) control vs permissive and 2) permissive vs non-permissive conditions.**

S2 File **List of gene identifiers used to generate each TMT figure.**

Table S1 ***SBF-EM analysis of different cell cycle stages identified a high proportion of cells containing disordered kDNA following TbFUT1 KO***

Figure S1: ***Loss of TbFUT1 causes accumulation of dyskinetoplastic cells and impairs ΨΔm generation***

Figure S2: ***Detection of fragmented kDNA by overexposure of DAPI stained TbFUT1 cKO mutants grown -Tet***

Figure S3: ***Validation of single allele (TbFUT1 cKO/F1-$\gamma^{WT/L262P}$) and double allele (TbFUT1 cKO/F1-$\gamma^{L262P/L262P}$) point mutations in $F_1$-γ mutants***

Figure S4 ***Comparisons of steady state protein levels between wild-type and TbFUT1 cKO mutant grown in + Tet conditions to induce TbFUT1 overexpression***

Figure S5 ***Gel slices excised to perform protein identification of $F_oF_1$-ATP synthase complex subunits from wild-type and TbFUT1 cKO mutants grown ±Tetracycline for 3 days***

Figure S6 ***F1-γ L262P mutants depleted of TbFUT1 exhibit similar levels of transcripts ± Tet as parental TbFUT1 cKO cells***

| Cell type | +Tet dataset | | | -Tet dataset | | |
|---|---|---|---|---|---|---|
| | Total number of cells counted for each stage | Number with ”disordered kDNA” | % Cells with “disordered kDNA” | Total number of cells counted for each stage | Number with ”disordered kDNA” | % Cells with “disordered kDNA” |
| G1 | 8 | 2 | **25** | 27 | 22 | **81.5** |
| S-phase - mitosis | 9 | 0 | **0** | 14 | 12 | **85.7** |
| Post-mitotic | 1 | 0 | **0** | 4 | 2 | **50** |
| Cytokinesis | 0 | 0 | **0** | 1 | 0 | **0** |
| Total | 18 | 2 | **11.1** | 46 | 36 | **78** |

Table S1 ***SBF-EM analysis of different cell cycle stages identified a high proportion of cells containing disordered kDNA following TbFUT1 KO.*** TbFUT1 cKO cells grown ± Tetracycline for 77 h were used for cell counting to quantify kDNA disorder. Numbers of cells counted and the corresponding % of total population are indicated.

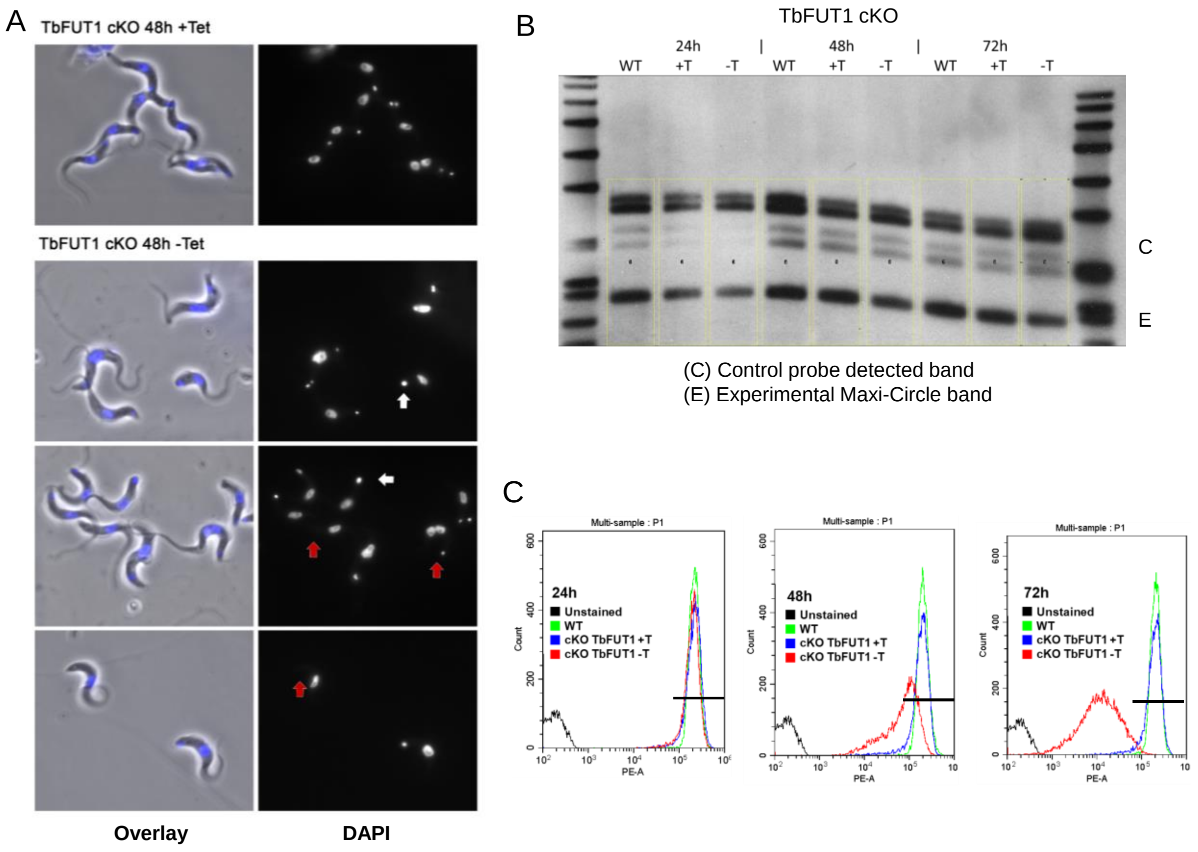


Figure S1: ***Loss of TbFUT1 causes accumulation of dyskinetoplastic cells and impairs ΨΔm generation***
A. Representative images showing TbFUT1 KO mutants imaged 2 days following growth in the presence (+Tet) or absence (-Tet) of tetracycline. DNA stained by DAPI in blue in the overlay panel. Red arrows indicate cells lacking any detectable kDNA, white arrows indicate cells with kDNA of increased size. B. Southern blot detection of Maxicircle kDNA (band E) harvested from cells 24, 48 and 72 hours following TbFUT1 KO (-Tet). Quantification of Maxicircle DNA relative to the loading control (C) was calculated using ImageJ gel peak analysis software. The normalised expression values were used to compare band intensity between TbFUT1 cKO cells ± Tet. C. Flow cytometry analysis of live WT and TbFUT1 cKO mutants cells treated with TMRE after growth in the presence (+Tet) or absence (-Tet) of tetracycline for 24, 48 and 72 hours. A non-stained WT control was included as a negative control and a TMRE positive gate drawn (black line) based on wild type fluorescence levels.

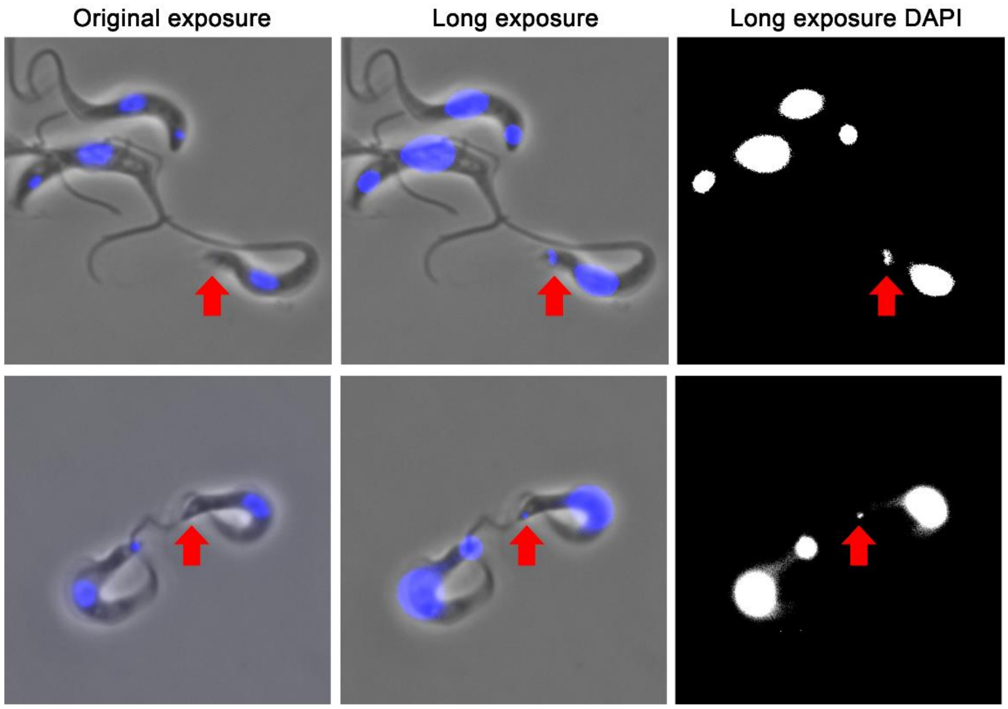


Figure S2: ***Detection of fragmented kDNA by overexposure of DAPI stained TbFUT1 cKO mutants grown -Tet.*** Low levels of fluorescent signal emanating from the kDNA region were observed in TbFUT1 KO mutants.

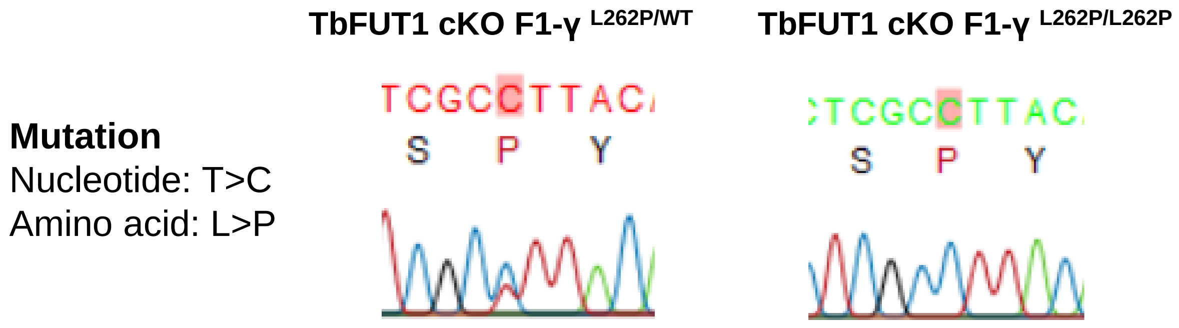


Figure S3: ***Validation of single allele (TbFUT1 cKO/F1-γ$^{WT/L262P}$) and double allele (TbFUT1 cKO/F1-γ$^{L262P/L262P}$) point mutations in F$_1$-γ mutants***

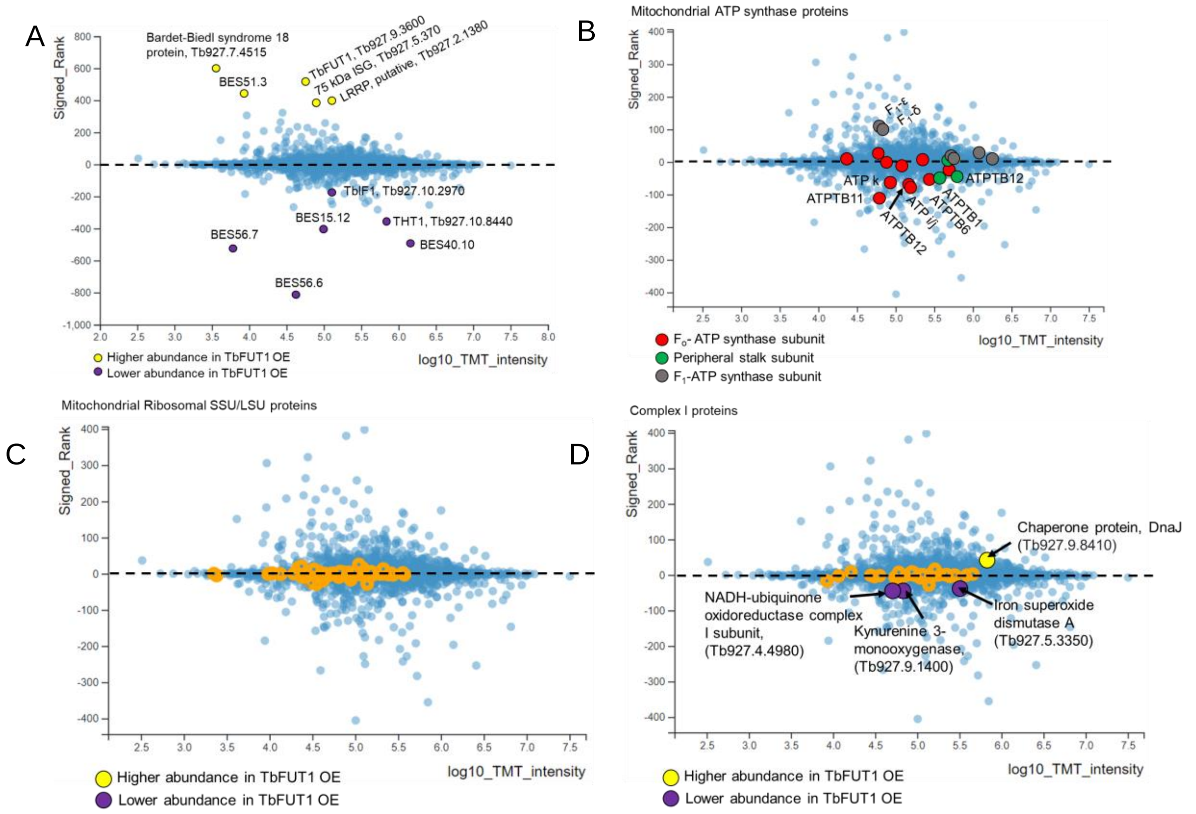


Figure S4 ***Comparisons of steady state protein levels between wild-type and TbFUT1 cKO mutant grown in + Tet conditions to induce TbFUT1 overexpression*** A. Panel indicating the increase (yellow circles) or decrease (purple circles) in a cohort of proteins, including bloodstream expression site (BES) associated proteins following TbFUT1 overexpression (OE). B. The steady state levels of $F_o$-subcomplex proteins (red circles) and peripheral stalk components (green circles) are lower in TbFUT1 cKO cells grown +Tet relative to WT. In contrast, $F_1$-subcomplex subunits (grey circles) are stable or more abundant in TbFUT1 OE cells. C. Subunits of the mitochondrial ribosomal LSU and D. Complex I remained stable following TbFUT1 OE.

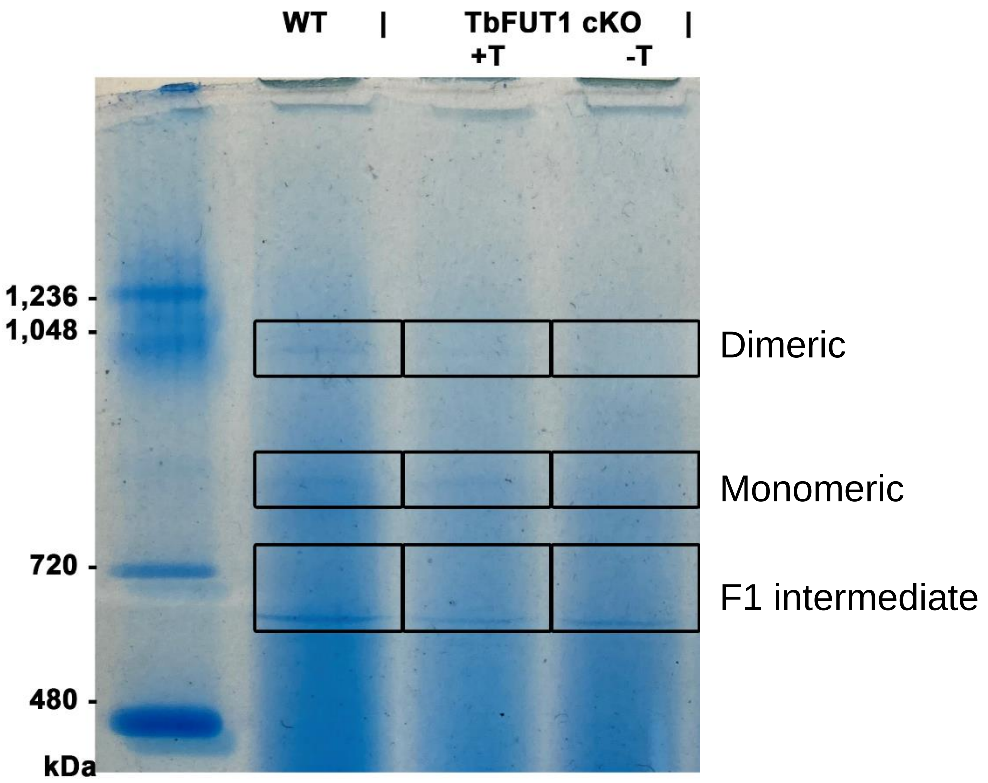


Figure S5 ***Gel slices excised to perform protein identification of $F_oF_1$-ATP synthase complex subunits from wild-type and TbFUT1 cKO mutants grown ±Tetracycline for 3 days.*** Native PAGE gel was fixed by Quick Coomassie and gel slices corresponding to $F_1$ intermediates, monomeric and dimeric $F_oF_1$-ATP synthase complexes were excised and subjected to LC-MS/MS analysis.

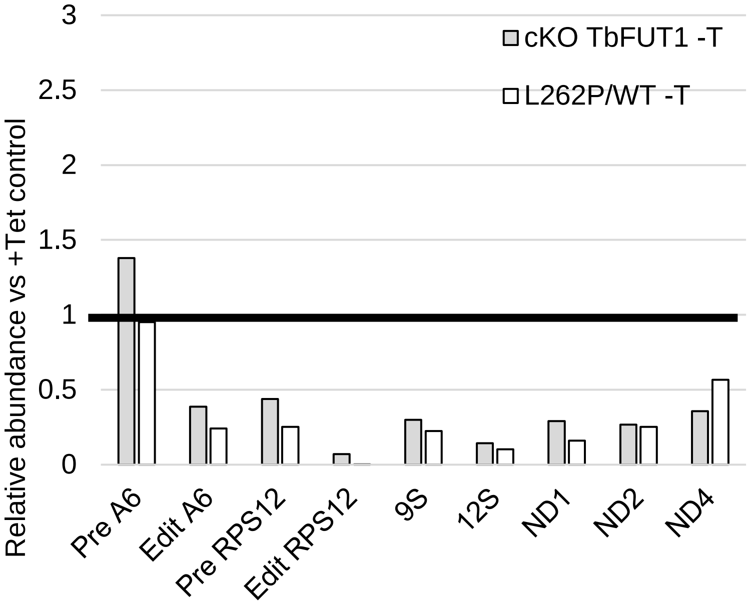


Figure S6 ***F1-γ L262P mutants depleted of TbFUT1 exhibit similar levels of transcripts ± Tet as parental TbFUT1 cKO cells.*** TbFUT1 cKO and TbFUT1 cKO/F1-$\gamma^{WT/L262P}$ mutants. Cells were grown ±Tet for 48 h and RNA harvested and analysed by RT-qPCR to detect kDNA encoded transcript levels. Relative abundance of transcripts are shown by normalising against +Tet (TbFUT1 expressing) controls.
